# Extracellular matrix regulates morphogenesis and function of ciliated sensory organs in *Caenorhabditis elegans*

**DOI:** 10.1101/376152

**Authors:** Deanna M. De Vore, Karla M. Knobel, Ken C.Q. Nguyen, David H. Hall, Maureen M. Barr

## Abstract

Cilia and extracellular vesicles (EVs) are signaling organelles that play important roles in human health and disease. In *C. elegans* and mammals, the Autosomal Dominant Polycystic Kidney Disease (ADPKD) gene products polycystin-1 and polycystin-2 localize to both cilia and EVs, act in the same genetic pathway, and function in a sensory capacity, suggesting ancient conservation. Hence, the nematode offers an excellent system in which to address central questions regarding the biology of cilia, EVs, and the polycystins. We discovered an unexpected role of the *mec-1, mec-5*, and *mec-9* genes encoding extracellular matrix (ECM) components. We determined that these ECM encoding genes regulate polycystin localization and function, ciliary EV release, cilia length, dendritic morphology, and neuron-glia interactions. Abnormal ECM and fibrosis are observed in ciliopathies such as ADPKD, nephronophthisis, and Bardet-Biedl Syndrome. Our studies reveal multifaceted roles for ECM proteins in the ciliated nervous system of the worm and provide a powerful new *in vivo* model to study the relationship between ECM, the polycystins, and ciliopathies.

## INTRODUCTION

Cilia are antenna-like structures that project from many eukaryotic cells (Wood and Rosenbaum 2015). Cilia play essential roles in human development and health, with ciliary defects resulting in syndromic ciliopathies (Reiter and Leroux 2017). Cilia are endowed with receptors, channels, and signaling components, enabling cilia to act as cellular sensors. In addition to their sensory abilities, cilia may transmit signals via submicroscopic extracellular vesicles (Wang and Barr 2018; Wood and Rosenbaum 2015). The mechanisms that enable a cilium to simultaneously send and receive information remain mysterious.

Cilia and the extracellular matrix (ECM) share an intimate association, with cilia projecting into and being surrounded by ECM (Seeger-Nukpezah and Golemis 2012). ECM is made up of a network of interacting proteins that surround and support cells for adhesive, structural and signaling functions (Hynes 2009). ECM is necessary for tissue morphogenesis and homeostasis throughout the lifespan of an organism and dynamically interacts with and regulates body systems, organs, and tissues. In the brain and nervous system, ECM is important for neuronal development, anatomy, and synaptic transmission. Dysregulation of ECM contributes to pathological conditions such as invasive cancer and fibrosis (Bonnans *et al*. 2014). Fibrosis and abnormal ECM are observed in ciliopathies such as autosomal dominant polycystic kidney disease (ADPKD), nephronophthisis, and Bardet-Biedl Syndrome (Song *et al*. 2017).

ADPKD is a genetic disorder characterized by the presence of fluid filled cysts which form on and in the kidney epithelia lining renal tubules that replace normally functioning renal tissue and result in enlarged, polycystic kidneys and end stage renal failure (Ghata and Cowley 2017; Ong and Harris 2015). ADPKD affects 1 in ~500 persons regardless of race or gender and can be caused by a mutation in either of the polycystin encoding genes PKD1 or PKD2. The large extracellular domain of polycystin-1 extends into the ECM and contains domains that may mediate protein-protein or protein-ECM interactions. PKD2 has an often-mutated extracellular domain that is necessary for polycystin channel assembly, stimulation, and gating (Shen *et al*. 2016). These two polycystins can be cleaved and have been found to act individually and synergistically, performing myriad functions including acting as ion channels and regulation of ECM components (Hanaoka *et al*. 2000; Mangos *et al*. 2010). Polycystins and ECM seem to mutually regulate each other, suggesting dynamic feedback.

The polycystins localize to cilia and extracellular vesicles, and this subcellular localization is evolutionarily conserved and observed in *Chlamydomonas, C. elegans*, and mammals (Hogan *et al*. 2009; O’Hagan *et al*. 2014; Semmo *et al*. 2014; Wang *et al*. 2014; Wood *et al*. 2013; Wood and Rosenbaum 2015). EVs carry many ECM proteins such as fibronectin and laminin, which provide communication necessary for altering ECM composition, signaling between ECM and the cells it surrounds, tumor proliferation, and inflammation (Rilla *et al*. 2017). In *Chlamydomonas*, ciliary EVs carry a proteolytic enzyme that degrades ECM required for hatching (Wood *et al*. 2013). In *C. elegans*, ciliary proteins such as polycystins are EV cargo that function in animal-to-animal communication (Wang *et al*. 2014).

The *C. elegans* polycystins LOV-1 and PKD-2 localize to cilia of male-specific sensory neurons (Barr *et al*. 2001; Barr and Sternberg 1999). We previously performed a forward genetic screen for regulators of PKD-2::GFP ciliary localization (Bae *et al*. 2008). Here we identify a mutation in the collagen gene *mec-5* that produced a PKD-2::GFP ciliary localization defect and discovered new functions for the *mec-1, mec-5*, and *mec-9* ECM genes previously implicated in the function of non-ciliated touch receptor neurons (Katta *et al*. 2015). MEC-1 and MEC-9 contain EGF/Kunitz domains and MEC-5 is a Type IV collagen (Emtage *et al*. 2004). MEC-1, MEC-5, and MEC-9 ECM proteins form the mantle surrounding non-ciliated touch receptor neurons, mediate touch neuron attachment to the hypodermal skin of the worm, and regulate the localization of the mechanosensitive DEG/ENaC (degenerin/sodium epithelial channel) complex MEC-4 and MEC-10 (Du *et al*. 1996; Gu *et al*. 1996).

Here we demonstrate that *mec-1, mec-5*, and *mec-9* are required for polycystin protein localization to the sensory cilia and discovered that proteins in the extracellular matrix regulate the movement and activity of proteins inside the cell. We find that these ECM components also regulate polycystin-mediated male mating behaviors, control neuron-glia interactions important for ciliary and dendritic integrity, and modulate the shedding and release of ciliary extracellular vesicles. While the polycystins have been implicated in sensing and regulating collagen in zebrafish models, roles for ECM proteins in regulating ciliary integrity, ciliary polycystin localization, and ciliary function have not been previously appreciated.

## MATERIALS AND METHODS

### Culture of *C. elegans* nematodes

Nematodes were maintained using standard conditions (Brenner 1974). Males and hermaphrodites were isolated at L4 stage d24hrs prior to experiments and kept at 20-22°C overnight. In *C. elegans*, the predominant sex is hermaphrodite and males spontaneously arise only rarely (less than 1%). Therefore, in all experiments in which males were tested, we used animals in either the *him-5(e1490)* or *him-8(e1494)* background. These backgrounds were considered wild type. *him-5(e1490)* and *him-8(e1494)* males exhibit normal mating behaviors and are used as wild-type controls for mating assays.

### General molecular Biology

PCR amplification was used for genotyping and building transgenic constructs using the following templates: *C. elegans* genomic DNA, cDNA, or prebuilt constructs. High fidelity LA Taq (TaKaRa Bio Inc., Otsu, Shiga, Japan) or Phusion High Fidelity DNA Polymerase (Thermo Fisher Scientific, Vantaa, Finland) were used for amplification of DNA for constructs. Sequencing reactions were performed by Genewiz, (South Plainfield, NJ, USA). DNA and protein sequence analysis BLAST was used for identification of gene orthologs in *C. elegans*. Human and nematode protein sequence information was provided by NCBI smartBLAST, and *C. elegans* gene and protein sequence information was also provided by WormBase. Serial Analysis of Gene Expression (SAGE) data provided by WormBase (Release WS221). Whole genome sequencing was performed and analyzed by Richard Poole using CloudMap. ApE 1.17 was used for sequence manipulation.

### Imaging

Nematodes were anaesthetized with 10 mM levamisole and mounted on agar pads for imaging at room temperature. Epifluorescence images were acquired using a Zeiss Axioplan2 microscope with 10x, 63x (NA 1.4), and 100x (NA 1.4) oil-immersion objectives with a Photometrics Cascade 512B CCD camera using Metamorph software (www.moleculardevices.com) or Zeiss Axio Imager.D1m microscope using a 63x and 100x objective with a Q imaging Regtiga-SRV camera. Optical Z-stack projections were stored as TIFF files and manipulated using ImageJ and Adobe Illustrator. Scale bars are 10 microns for head and tail images. EM scale bars are 200 or 500 nm as marked in image.

### Transmission Electron Microscopy

*mec-9(ok2853)* and wild-type young adult males were fixed using high-pressure freeze fixation and freeze substitution in 2% OsO4 + 2% water in acetone as the primary fixative (Weimer 2006). Samples were slowly freeze substituted in an RMC freeze substitution device, before infiltration with Embed-812 plastic resin. For TEM, serial sections (70-75 nm thickness) of fixed animals were collected on copper slot grids coated with formvar and evaporated carbon and stained with 4% uranyl acetate in 70% methanol, followed by washing and incubating with aqueous lead citrate. Images were captured on a Philips CM10 transmission electron microscope at 80kV with a Morada 11-megapixel TEM CCD camera driven by iTEM software (Olympus Soft Imaging Solutions). Images were analyzed using ImageJ (FIJI) and manipulated with Adobe Illustrator.

### Extracellular vesicle release

EV release was analyzed by counting all EVs from one-day old young adult males as described by Silva et al 2017. Individual animals were mounted on agar into four quadrants of agar slide. Free and newly released EVs that float to the cover slip were counted and reported as “number of EVs released.”

### Transmission Electron Microscopy EV measurements

Using ImageJ (FIJI) software, WT and mutant electron micrographs were compared at and around the CEM transition zone. Qualitative examination was performed. EV diameter measurements were performed by drawing a line across the widest diameter of an EV and taking a measurement via FIJI measurement tool reporting diameter in nm.

### Transmission Electron Microscopy cilia length measurements

Electron micrographs were stacked using the TrakEM2 component of ImageJ(FIJI) software. The transition zone was identified by Y-links and used as the bottom-most measurement. The ciliary tip was used as the top measurement. The Z stack position was multiplied by the thickness of the cut (~70-75 nm) and length was reported in μm.

### PKD-2 antibody generation and staining

Animals were staged young adults and washed off plates with M9. Antibodies were prepared using Abmart protocols and staining against PKD-2 was prepared and performed using a Finney Ruvkun protocol (Bettinger *et al*. 1996) [Wormatlas.org]. The monoclonal PKD-2 primary antibody was created by Abmart against the intracellular N-terminal domain (amino acids: DERWANPPQPVA) and an intracellular C-terminal domain (amino acids: KRGKRPDAPGED). The secondary antibody was α mouse Alexa Fluor ^®^ 568 donkey anti-mouse IgG (H + L) (2 mg/ml) by Invitrogen TM. The following dilutions and incubation times were used: primary antibodies 1:200 overnight (18-24 hours); secondary antibody 1: 1000 for two hours.

### Response behavior assay

Control Strains used: CB1490: *him-5(e1490)*, CB1489: *him-8(e1489)*, PT9: *pkd-2(sy606) him-5(e1490)*, and CB169: *unc-31(e169)IV*. L4 larval males were moved to a fresh plate approximately 24 hours before mating. *unc-31* mutant hermaphrodites were also picked as L4 larvae ~24 hours before experiments. Male mating assays were conducted on a fresh NGM agar plate with a small lawn of *E. coli* (OP50) containing 25 young-adult *unc-31* hermaphrodites. One, two, or three males were placed in the center of the lawn and observed for four minutes. When a male began scanning a hermaphrodite and the male tail maintained contact with a mate for at least ten seconds, a response was scored and that male was removed from the test plate.

### Location of vulva assay

was performed as described (Barr and Sternberg 1999). Location of vulva efficiency is calculated by successful vulva location divided by the total number of vulva encounters for each male. Total time measured was four minutes.

### Male leaving assay

was performed as described (Barrios *et al*. 2008). L4 males were picked and isolated from hermaphrodites on plates, then assayed 24 hours later in 20 μl food on a 9 cm diameter plate. Animals are positive for leaving when males exhibit tracks that approach within 1 cm of the edge of the plate. Time points scored were 2, 5, 8, and 24 hours after the males were placed on the spot of food. A minimum of 20 animals per strain and three replicates were scored for each genotype assayed. Statistical significance was determined by R software.

### Dye filling assays

Standard dye-filling assays (Perkins *et al*. 1986) were performed using Dil (Invitrogen). The number of amphid and phasmid cell bodies were counted, and the results reported as number of neurons out of 12 (amphid) or 4 (phasmid) that fill with DiI.

### Dendritic trafficking velocity measurements

To directly measure in vivo velocity of PKD-2::GFP in CEM dendrites, we acquired time lapse image stacks, which were later converted to kymographs using the KymographClear V2.0 plugin in FIJI. Motile particles were automatically and indiscriminately detected and traced, and velocities analyzed with KymographDirect (Mangeol *et al*. 2016). We observed a reduction of overall observed velocities compared to our previous publication (Bae *et al*. 2006), likely due to differences between automatic versus manual particle tracing. This reduction does not affect our analysis or overall conclusions, as our model is not based on absolute velocities, but rather on their relative changes.

### Ciliary localization and fluorescence intensity

Ciliary localization was performed by blind assay of a stack of images collected into a maximum intensity image using the “Z project” function in ImageJ software. Animals were scored as ciliary localization defective (Cil) if excess or misplaced PKD-2::GFP or endogenous PKD-2 (detected by α-PKD-2 antibodies) was detected (excess in cilium, ciliary base, or dendrite). Fluorescence intensity was measured using ImageJ by drawing a range of interest and using measurement tool. All measurements have background subtracted to create final value.

### Strains used: (transgenic lines created by Knudra)

CB1066 *mec-1(e1066)V*
CB1292 *mec-1(e1292)V*
CB1490 *him-5(e1490)*
CB1494 *mec-9(e1494)V*
CB1503 *mec-5(e1503)X*
CB169 *unc-31(e169)IV*
COP1472 *knuEx206[pNU1416-mec-9Sp:: GFP::tbb-2utr, unc-119(+)];unc-119(ed3)lll*
COP1473 *knuEx207[pNU1415-mec-9Lp::GFP::tbb-2utr,unc-119(+)];unc-119(ed3)III*
KU25 *pmk-1(km25)IV*
PT3296 *mec-9(ok2853)pmk-1(km25)IV;myIs4 him-5(e1490) V*
PT3168 *him-8(e1489) myIs1[PKD-2::GFP + cc::GFP] IV*
PT277 *unc-119(ed3)III; him-5(e1490) V*
PT1213 *myIs4 him-5(e1490) V; mec-5(my2) X*
PT1852 *pha-1(e2123) III; him-5(e1490) V; Ex [LOV-1::GFP1]*
PT2434 *dyf-1 (m335); myIs1; him-5*
PT2679 *him-5(e1490)V;myIs23[Pcil-7::gCIL-7::GFP_3’UTR+ccRFP]*
PT2962 *him-8(e1489) myIs1; mec-1(e1066) V*
PT2963 *him-8(e1489) myIs1; mec-1(e1336) V*
PT2964 *him-8(e1489) myIs1; mec-1(e1292) V*
PT2965 *him-8(e1489) myIs1; mec-9(e1494) V*
PT2966 *him-8(e1489) myIs1; mec-9(ok2853) V*
PT2967 *him-5(e1490) myIs4[PKD-2::GFP + cc::GFP] V; mec-5(e1790) X*
PT2968 *him-5(e1490) myIs4; mec-5(e1503) X*
PT2969 *uIs31[MEC-17:: GFP] III; him-5(e1490) myIs4[PKD-2:: GFP + cc::GFP] V; mec-5(u444) X*
PT3038 *unc-31(e169)IV; him-5(e1490), myIs4 V*
PT3203 *mec-9(ok2853); pha-1; him-5(e1490); syEx301[pBx+LOV-1::GFP1]*
PT3213 *mec-9(ok2853)V, him-5(e1490)V;myIs23[pcil-7::gCIL-7:: GFP:: 3’UTR+ccRFP]*
PT443 *myIs1 pkd-2(sy606) IV; him-5(e1490) V*
PT621 *myIs4 him-5(e1490) V*
RB2140 *mec-9(ok2853) V*

Strains and plasmids are available upon request. The authors affirm that all data necessary for confirming the conclusions of the article are present within the article, figures, and tables.

## RESULTS

### *mec-1, mec-5*, and *mec-9* regulate PKD-2::GFP localization

The *C. elegans* polycystins LOV-1 and PKD-2 localize to cilia and cell bodies of cephalic male-specific (CEM), left and right B-type ray neurons in the tail (RnB; n=neuron 1-9, but not 6), and Hook B (HOB) neurons. (Figure 1 A-B, Supplemental Figure 1 A-B) The polycystins and these male specific neurons are necessary for male mating behaviors (Barr *et al*. 2001; Barr and Sternberg 1999; Liu and Sternberg 1995; Srinivasan *et al*. 2008). We previously performed a forward genetic screen for genes necessary in PKD-2::GFP localization and identified mutants defective in PKD-2 ciliary receptor localization (the Cil phenotype) (Bae *et al*. 2008). The *cil-2(my2)* mutant displays abnormally high PKD-2::GFP levels in the CEM cilia and ciliary base and also has abnormal extracellular, extradendritic PKD-2::GFP accumulation along CEM dendrites (Bae *et al*. 2008). *cil-2(my2)* hermaphrodites also displayed temperature sensitive sterility phenotype, that is linked to the Cil phenotype. The extradendritic PKD-2::GFP accumulation and sterility phenotypes are unique to the *cil-2(my2)* mutant and not the other ten Cil mutants. We used genetic mapping, whole genome sequencing, complementation testing, and rescue experiments of the Cil phenotype to map the *my2* lesion to *mec-5*.

**Figure 1.**
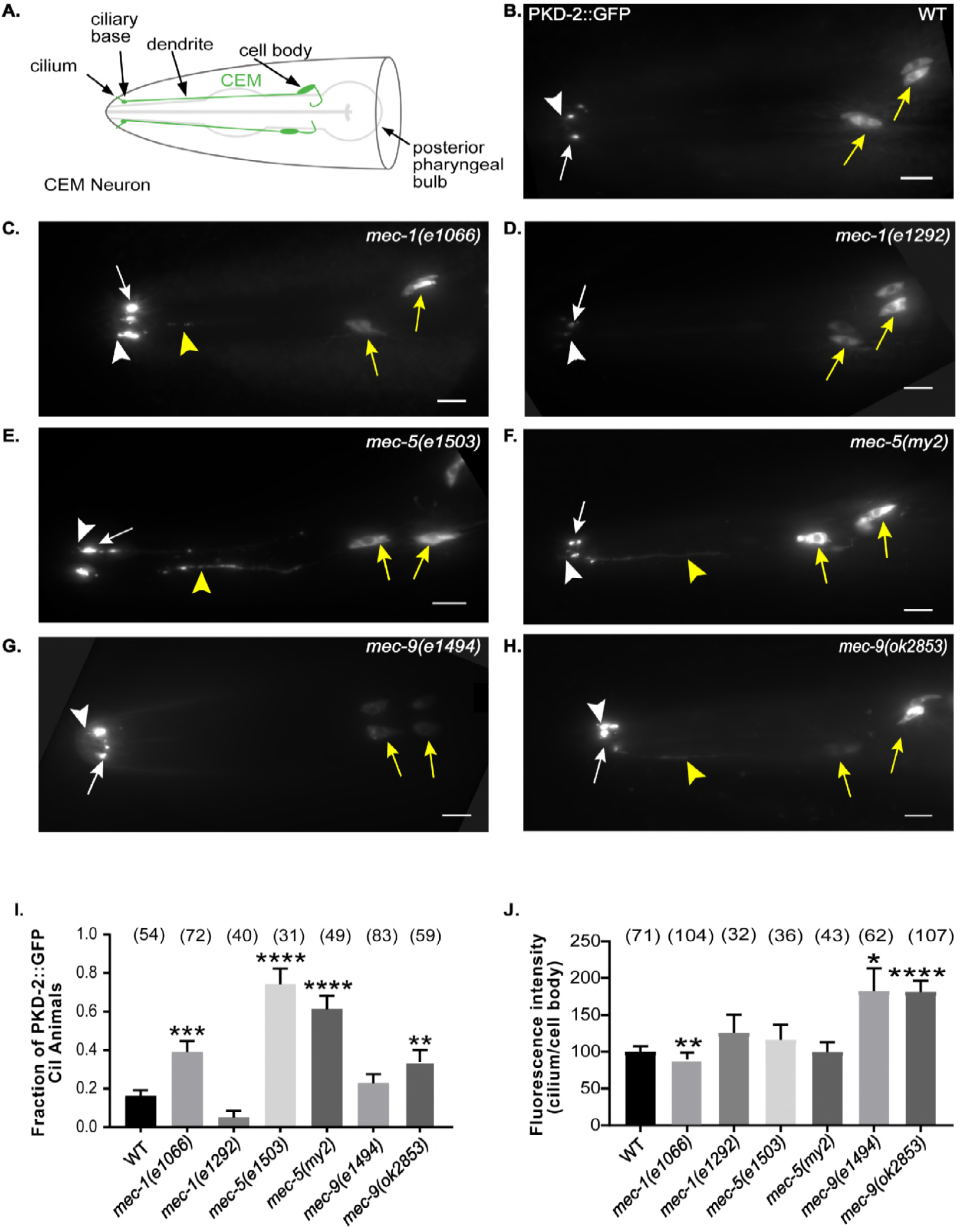
*mec-1, mec-5*, and *mec-9* regulate PKD-2::GFP localization and abundance. (B-H) Images were compiled from a 630x maximum intensity Z-series obtained by epifluorescence microscopy. Scale bar is 10 μm. White arrow, ciliary base; white arrow head, cilium; yellow arrow, cell body; yellow arrowhead, dendrite. (A) Schematic of WT cephalic male neurons in the head of a male (green). (B) A lateral view of WT PKD-2::GFP (translational reporter). PKD-2::GFP localized to CEM cell bodies and cilia. (C-D) Lateral view of PKD-2::GFP in *mec-1* mutant males. (C) Increased PKD-2::GFP at *mec-1(e1066)* CEM cilia and ciliary base. (D) *mec-1(e1292)* was indistinguishable from WT. (E-F) *mec-5* PKD-2::GFP ciliary localization. Extra-dendritic and increased ciliary PKD-2::GFP observed in *mec-5(e1503)* and *mec-5(my2)* male CEMs. (G-H.) PKD-2 localization in *mec-9* mutant males. (G) Increased PKD-2::GFP at *mec-9(e1494)* CEM ciliary base. (H) Extra-dendritic and increased PKD-2::GFP at *mec-9(ok2853)* CEM ciliary base. (I) Graph of the fraction of PKD-2::GFP mislocalization in CEM neurons of wild type and mutant males via blind examination. An animal was scored as “Cil” (ciliary localization defective) if CEM cilia, ciliary base, and/or dendrites in male worms had abnormally increased or mislocalized PKD-2::GFP. Values were reported as fraction of animals that are Cil. Significance was determined by Kruskal Wallace test with Dunn’s multiple comparisons test performed to compare groups. **p<0.01, ***p<0.001, ****p<0.0001. In WT, we observed occasional Cil animals (~16% of males). *mec-1, mec-5* and *mec-9* all had mutant alleles that mislocalized PKD-2::GFP. (J) Ratio (cilium/cell body) of maximum intensity showed that PKD-2::GFP abundance in CEM cilia is increased in comparison with the cell bodies only in *mec-9* ECM gene mutants; however, *mec-1* and *mec-5* alleles also affected PKD-2 abundance (Supplemental Table 1). Background measurements were subtracted from cilium and cell body values for standardization of images and we expressed the measurements in ratio of cilia to cell body FI. Significance was measured by Kruskal-Wallace test, comparisons made using Dunn’s multiple comparisons. *p<0.05; **p<0.01; ***p<0.001; **** p<0.0001. WT values were normalized to 100. The *mec-9* mutants had a brighter maximum FI (~1.75x) than WT (Figure 1J).

Expression of the wild-type (WT) *mec-5* genomic region using either the *mec-5* promoter or muscle-specific *myo-3* promoter rescued *cil-2(my2)* defects (data not shown), confirming that *mec-5* was mutated in the *cil-2(my2)* mutant and that *mec-5* acted non-cell autonomously to regulate PKD-2::GFP ciliary localization in male-specific neurons. *mec-5* encodes a collagen protein that is produced and secreted by hypodermal cells to anchor the degenerin complex in touch receptor neurons to the extracellular matrix (ECM) (Du *et al*. 1996). In these non-ciliated neurons, *mec-5* and the ECM encoding genes *mec-1* and *mec-9* act in concert (Du *et al*. 1996). We therefore determined whether MEC-1, MEC-5, and MEC-9 also regulated PKD-2::GFP ciliary localization.

We characterized PKD-2::GFP ciliary localization in two mutant alleles each of *mec-1* (*e1066* and *e1292*), *mec-5* (*e1503* and *my2*), and *mec-9* (*e1494* and *ok2853*) (Figure 1C-H). First, we visualized ciliary localization in CEM neurons of WT and mutant males via blind examination. A male was scored as Cil if CEM cilia, ciliary base, and dendrites in male worms had abnormally increased or mislocalized PKD-2::GFP. Values were reported as percent of animals that are Cil (Figure 1I). In WT, we saw occasional Cil animals (~16% of males). In contrast, *mec-5* males exhibited the most penetrant Cil phenotype: over 60% of *mec-5(my2)* animals and over 70% of *mec-5(e1503)* were Cil (p values < 0.0001). Also, *mec-9(ok2853)* animals had a significantly increased number of Cil males (p=0.008), but *mec-9(e1494)* males were statistically indistinguishable from WT. *mec-1(e1066)* males were also Cil (p= 0.0003), but *mec-1(e1292)* males were similar to WT.

In *mec-1(e1066), mec-5(e1503), mec-5(my2)*, and *mec-9(ok2853)* males, we observed abnormally increased PKD-2 localization in CEM cilia, ciliary bases, and occasionally dendrites. (Figures 1C, E, F, and H) The extradendritic accumulation in *mec-1(e1066), mec-5(e1503)* and *mec-9(ok2853)* was reminiscent of that observed in *mec-5(my2)*. The Cil defect was not observed in *mec-1(e1292)* and was only occasionally seen in *mec-9(e1494)*. (Figures 1D and G)

*mec-1* encodes multiple ECM protein isoforms with multiple disulfide-linked EGF and Kunitz-type protease inhibitor domains (Emtage *et al*. 2004). The *mec-1* alleles cause gene truncations: *e1292* results in truncation after Kunitz Domain 3 whereas *e1066* truncates after Kunitz Domain 6 (Emtage *et al*. 2004), both of which remove Kunitz domains but might leave EGF domains intact. *mec-9* encodes two protein isoforms of an ECM protein containing multiple EGF domains, multiple Kunitz domains, and a glutamic acid-rich region (Du *et al*. 1996). *mec-9* mutations perturb EGF domains: *e1494* is a point mutation in the first set of EGF domains and affects only the *mec-9* long *(mec-9L)* isoform. *ok2853* perturbs the second set of EGF domains via deletions and affects both short and long isoforms (Du *et al*. 1996). Using whole genome sequencing, we found that both *mec-5* mutants had lesions in the third intron (data not shown). We conclude that some but not all alleles of *mec-1, mec-5* and *mec-9* perturbed PKD-2::GFP localization.

We observed that cilia and ciliary bases appeared brighter in some but not all ECM mutants. We measured highest/maximum fluorescence intensity (FI) of PKD-2::GFP in CEM neuron cell bodies and cilia (including both cilium and ciliary base) to quantify PKD-2::GFP abundance. FI is a computed measurement of the pixels illuminated in a selected region of interest. *mec-9* mutants had a brighter maximum FI (~1.75x) than WT (Figure 2B): *mec-9(e1494)* p=0.011 and *mec-9(ok2853)* p<0.0001 (Figure 1J). *mec-1(e1066)* mutants were dimmer than WT: p=0.0088. We observed variability in CEM FI of ECM gene mutant cilia, dendrites, and cell bodies (Supplemental Table 1), with *mec-9(ok2853)* exhibiting the brightest overall FI. Similar to CEM neurons, we also observed variable ray neuron FI in ECM mutant tails (Supplemental Figure 1 and Supplemental Table 1), again with *mec-9(ok2853)* exhibiting the brightest overall FI. Our data shows that although *mec-1, mec-5*, and *mec-9* were all necessary for PKD-2 ciliary localization, *mec-9* also regulated PKD-2::GFP ciliary abundance.

**Figure 2.**
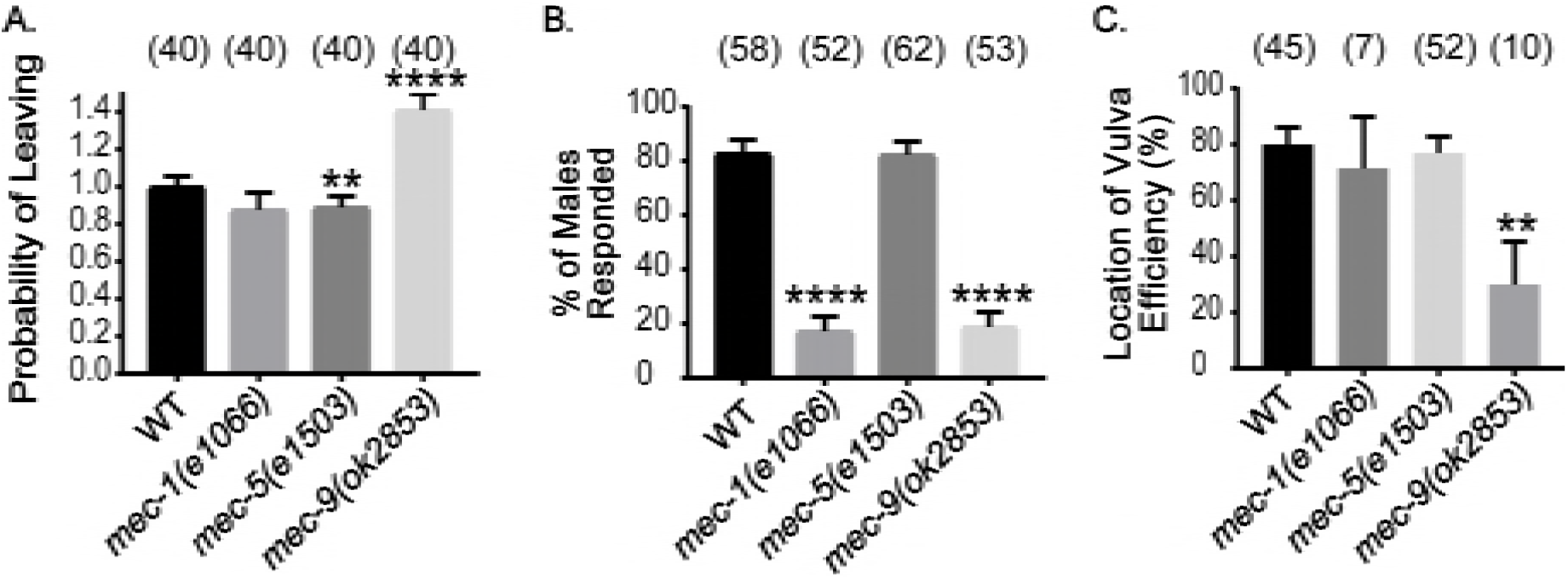
*mec-1, mec-5*, and *mec-9* are required for male mating behaviors. (A-C) *mec-1(e1066)* males were response defective, *mec-5(e1503)* males were leaving assay defective, *mec-9(ok2853)* males were defective in all three mating behaviors. (A) Leaving assay measured the probability of males leaving food to search for a mate. WT values normalized to 1. *mec-5 (e1503)* males were leaving assay defective when compared to WT. *mec-9(ok2853)* males left food more readily than WT. (B) *mec-1(e1066)* and *mec-9(ok2853)* males were defective in responding to hermaphrodite contact. (C) *mec-9(ok2853)* males were location of vulva defective. Significance determined by Kruskal-Wallis test (Mann-Whitney test for WT vs. *mec-5(e1503)* only); p values compared by Dunn’s multiple comparisons test. ** p<.01; **** p<0.0001.

To test for specificity of ECM components, we examined PKD-2::GFP localization in a hemicentin mutant. Hemicentin is an ECM component required for adhesion between tissues including touch neuron attachment to the epidermis and gonad (Vogel and Hedgecock 2001). Analysis of the hemicentin mutant *him-4(e1267)* revealed no differences in PKD-2::GFP ciliary localization or FI. We conclude that the *mec-1, mec-5*, and *mec-9* ECM genes; but not the *him-4* ECM gene, were necessary for polycystin localization and abundance.

We generated a monoclonal anti-PKD-2 antibody to visualize and measure endogenous PKD-2 localization in WT and ECM gene mutant males and observed similar Cil defects (Supplemental Figures 2 and 3). In WT, endogenous PKD-2 was limited to the cell bodies and cilia of CEM head neurons and ray RnB and hook HOB tail neurons (Supplemental Figure 3A and 3B). In *mec-9(ok2853)* males, endogenous PKD-2 mislocalized to dendrites and had increased abundance in cilia and cell bodies as shown in FI measurements (Supplemental Figure 3C and 3D). Endogenous PKD-2 localization was also abnormal in the ray cilia in *mec-1(e1066)* and *mec-5(e1503)* (data not shown) male tail but not as severely as *mec-9(ok2853)*. We conclude that all three ECM genes regulate PKD-2 localization with *mec-9* mutants displaying the most severe Cil defects.

The partner of Polycystin-2 (PKD-2) is Polycystin-1 (LOV-1). We therefore, examined LOV-1 localization in ECM gene mutants. In WT, LOV-1::GFP localizes to the CEM, RnB, and HOB cell bodies and cilia. In *mec-9(ok853)* mutants, we observed distal dendritic LOV-1::GFP mislocalization and increased ciliary fluorescence (Supp. Figure 4). We also observed significantly increased LOV-1::GFP FI in CEMs and RnBs of *mec-5(e1503)* males (data not shown). We conclude that ECM encoding genes *mec-1, mec-5*, and *mec-9* regulate polycystin localization in male-specific ciliated sensory neurons.

### *mec-1, mec-5*, and *mec-9* regulate male mating behaviors

*pkd-2, lov-1*, and the male-specific polycystin expressing neurons are required for the male mating behaviors of mate searching, response to hermaphrodite contact, and location of hermaphrodite vulva (O’Hagan *et al*. 2014). We therefore determined whether the three ECM genes were required for these male sensory behaviors.

*C. elegans* males leave a food source in search of a mate if no hermaphrodite is present (Lipton *et al*. 2004). *lov-1* and *pkd-2* mutant males do not leave food to search for a mate (Barrios *et al*. 2008). Similarly, *mec-5(e1503)* males were leaving defective (Figure 2A). In contrast, *mec-9(ok2853)* mutants left food more readily than WT animals (Figure 2A). Hyper-leaving behavior is associated with defects in male-specific and the shared inner labial type 2 IL2 ciliated neurons (Maguire *et al*. 2015), suggesting that the *mec-9* mutation may affect other neurons in addition to the polycystin-expressing cells.

When male ray neurons detect contact with a hermaphrodite, males initiate the response behavior by stopping forward locomotion, and initiating backing (Barr and Garcia 2006; Barr *et al*. 2018). *lov-1* and *pkd-2* mutant males are response defective. *mec-9(ok2853)* and *mec-1(e1066)* mutants were also defective in response to hermaphrodite contact, while *mec-5(my2)* mutant males displayed normal response behavior (Figure 2B) (Bae *et al*. 2006). After response, the male scans the hermaphrodite’s body for her vulva. *lov-1* and *pkd-2* mutants are location of vulva defective (Lov). Only *mec-9(ok2853)* males displayed the Lov phenotype (Figure 2C).

We previously showed that mate searching, response, and location of vulva do not require MEC-4 and MEC-10, the touch neuron specific degenerin epithelial sodium channel (DEG/ENaC) receptors (Barr and Sternberg 1999; Barrios *et al*. 2008). However, *mec-1, mec-5*, and *mec-9* ECM genes are required for these male-specific sensory behaviors. We conclude that *mec-1, mec-5* and *mec-9* ECMs genes act beyond DEG/ENaC localization and function in touch receptor neurons and are required more broadly for the function of other sensory neurons. Moreover, only *mec-9* was required for all examined male-specific behaviors suggesting a distinct role for *mec-9* in polycystin-expressing male specific neurons.

### *mec-9* long and short isoforms have distinct expression patterns: MEC-9S is expressed in ciliated sensory neurons

*mec-9* encodes two predicated proteins (Du *et al*. 1996). The *mec-9* long isoform encodes an 839-amino acid protein with five Kunitz protease inhibitor domains, seven EGF domains, and a glutamic acid rich/coiled-coiled region (Figure 3A) (Du *et al*. 1996). The *mec-9* short isoform encodes a 502-amino acid protein with two Kunitz domains, three EGF domains, and a glutamic acid rich/coiled-coiled region (Du *et al*. 1996) (Figure 3B). Both long and short isoforms encode an N-terminal signal sequence consistent with secreted ECM proteins. *mec-9(ok2853)* is a 408-base pair in-frame deletion that is predicted to remove two of the three EGF like domains in both isoforms (Consortium 2012) (Figure 3A).

Chalfie and colleagues showed *mec-9* long isoform expression in touch receptor neurons and *mec-9* short isoform expression in other neurons in the nerve ring and ventral cord in hermaphrodites (Du *et al*. 1996). We examined long and short isoform expression patterns in males and hermaphrodites using transcriptional reporters (Figure 3A). We find that *mec-9Lp::GFP* (long isoform transcriptional reporter) was expressed only in the shared touch receptor neurons (found in both males and hermaphrodites) and not male-specific neurons. However, *mec-9Sp::GFP* (short isoform transcriptional reporter) was expressed in polycystin-expressing CEM, RnB, and HOB male-specific as well as shared ciliated sensory neurons found in both males and hermaphrodites (Figures 3D and 3E).

**Figure 3.**
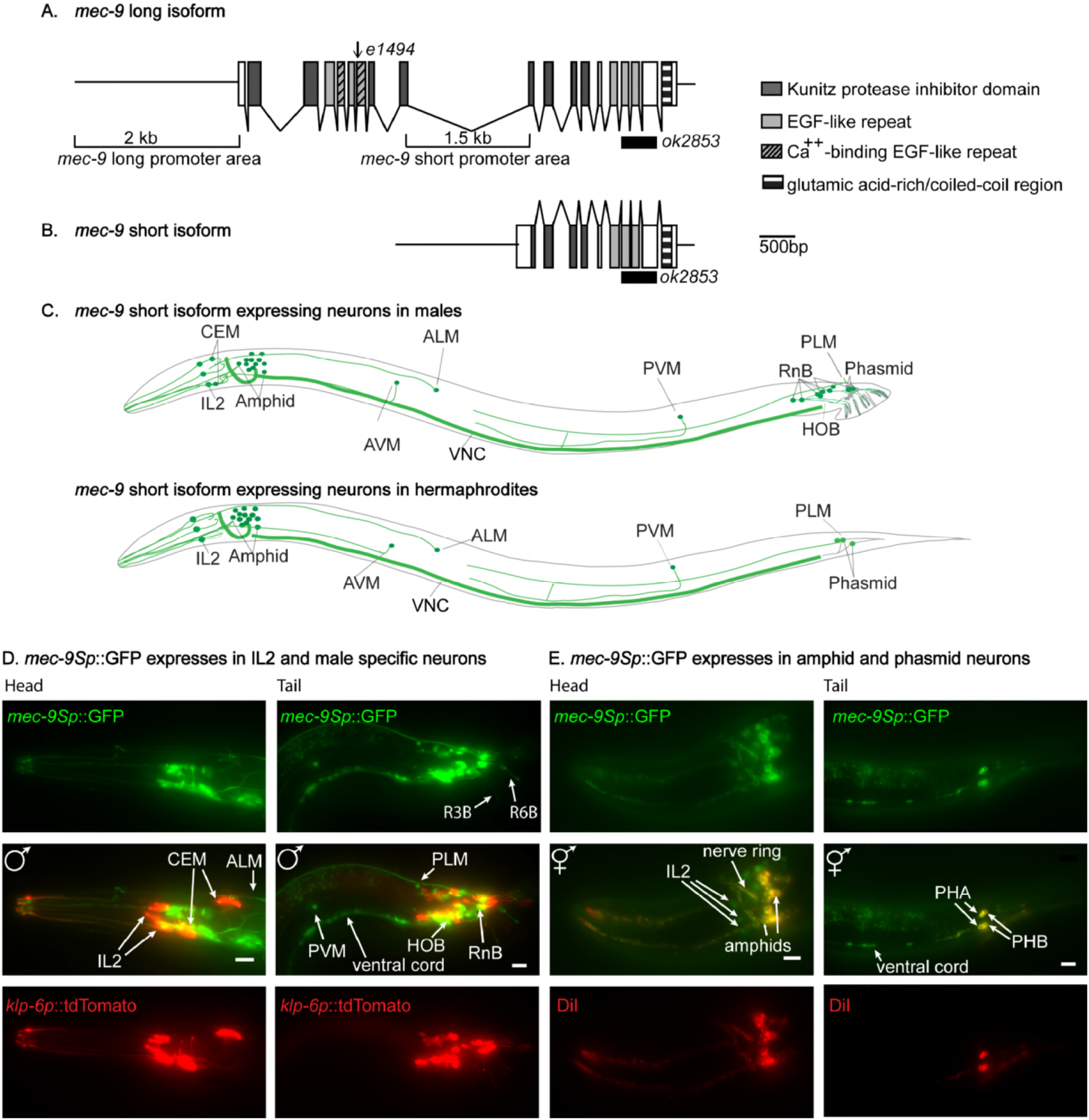
*mec-9Sp::GFP* is expressed in ciliated sensory neurons. (A and B) *mec-9* long and short transcripts. (A) *mec-9* long isoform encodes a secreted ECM protein that has 5 Kunitz domains, 7 EGF repeats, and a glutamic acid-rich/coiled-coil region (Du et al., 1996). *e1494* is a point mutation in the first set of EGF domains and *ok2853* perturbs the second set of EGF domains via base pair deletions. (B) *mec-9* short transcript is 502 aa (Du et al., 1996): the promoter for the *mec-9* short transcript located in *mec-9* long 9th intron. We used the 2kb upstream sequence with a C terminal GFP tag to construct *mec-9Lp::GFP* transcriptional reporter. We used the 1.5kb *mec-9* long intron 9 sequence with a C terminal GFP tag to construct *mec-9Sp::GFP* transcriptional reporter. (C) Ciliated sensory neurons expressed *mec-9Sp::GFP:* CEMs, IL2, amphids, RnB, HOB, phasmids. Male (top), hermaphrodite (bottom). Ventral nerve cord (non-ciliated neurons) expression also observed and shown. (D) *mec-9Sp::GFP* coexpressed with the kinesin-3 gene, *klp-6p::tdTomato* transcriptional reporter. *klp-6p::tdTomato* expressed in male-specific CEM and IL2 neurons in the head and RnB and HOB male-specific neurons in the male tail. Left column is lateral view of male head. *mec-* 9Sp::GFP expressed in CEMs and IL2 neurons. Right column is lateral view of male tail. *mec-9Sp::GFP* expressed in HOB and RnB neurons. (E) Some *mec-9Sp::GFP* expressing ciliated neurons filled with Dil. Dil is a lipophilic dye taken up by amphids and phasmids. Left column is lateral view of hermaphrodite head. *mec-9Sp::GFP* expresses in amphids in the hermaphrodite and male head. Right column is lateral view of hermaphrodite tail. *mec-9Sp::GFP* expressed in phasmids.

*mec-9Sp::GFP* was coexpressed with the kinesin-3 gene *klp-6* transcriptional reporter in the shared IL2 neurons and male-specific CEM neurons in the head and RnB and HOB neurons in the tail (Figure 3D). These 27 *klp-6* expressing neurons are called extracellular vesicle releasing neurons (EVNs) based on their ability to shed and release ciliary EVs (Wang et al., 2015). *mec-9Sp::GFP* was also expressed in the amphid neurons in the head and phasmid in the tail that take up the lipophilic fluorescent dye Dil (Figure 3E).

### MEC-9 regulates extracellular vesicle biogenesis and release

EVs are shed and released from ciliated IL2, CEM, and RnB neurons (Wang *et al*. 2014), all of which expressed *mec-9Sp::GFP*. To determine if *mec-9* regulates EV biogenesis, we counted PKD-2::GFP containing EVs that were shed and released into the local environment from the male head and tail. WT adult males released an average of 26 PKD-2::GFP labeled EVs from the CEM neurons compared to 78 EVs in *mec-9* mutants (p=0.0008): a threefold increase (Figure 4C). The EV hypersecretion phenotype of *mec-9* mutants contrasts other previously described EV hyposecretion mutants that are deficient in EV release (Maguire *et al*. 2015; O’Hagan *et al*. 2017; Silva *et al*. 2017; Wang *et al*. 2014).

**Figure 4.**
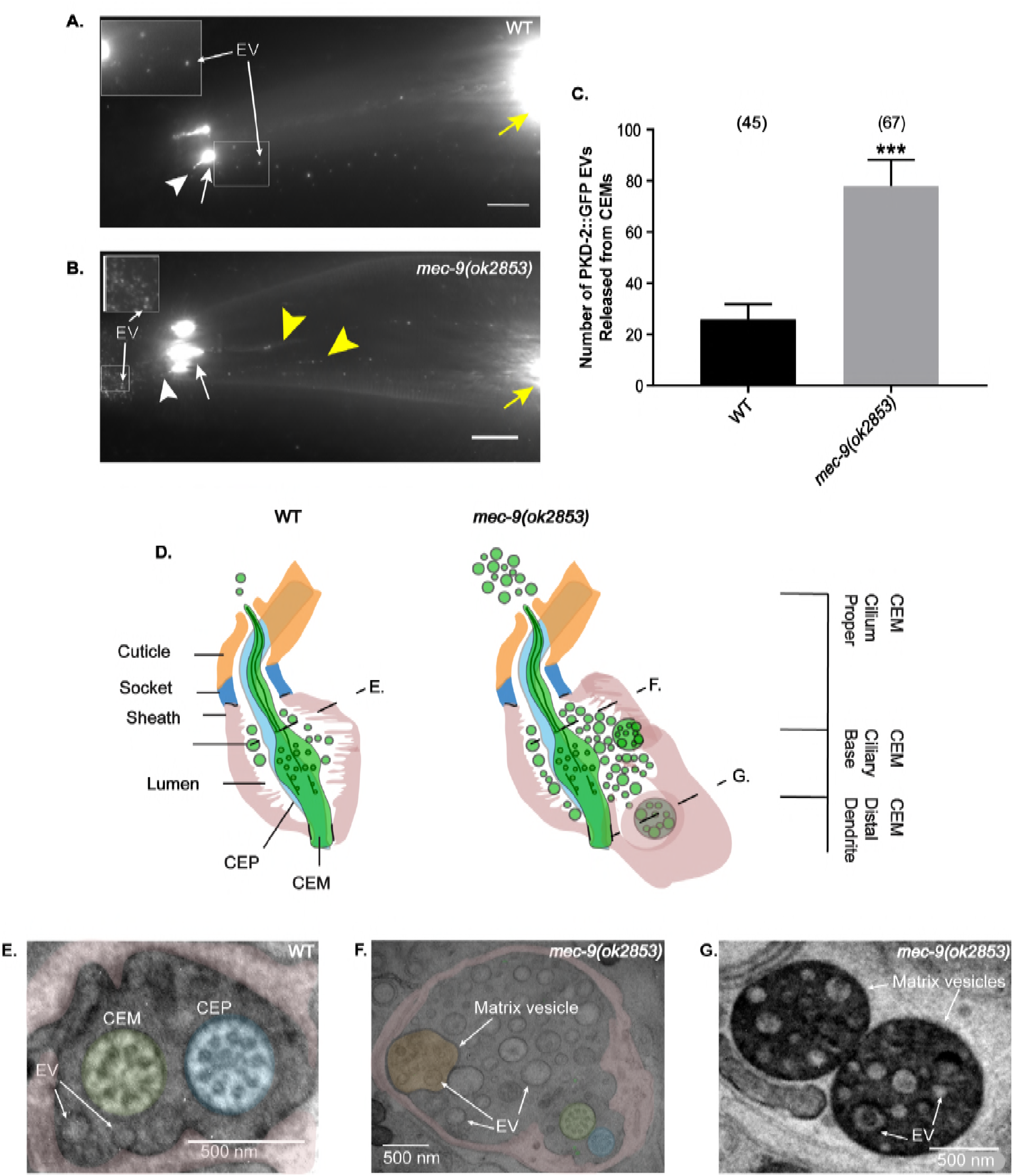
*mec-9(ok2853)* males produce excessive EVs and matrix-filled vesicles. (A-B) Head images of WT and *mec-9(ok2853)* mutant males show EVs were released. Image brightness was increased and inset was magnified to visualize EVs. Scale bars 10μm. White arrow, ciliary base; white arrow head, cilium; yellow arrow, cell body; yellow arrowhead, dendrite. (B) *mec-9(ok2853)* males produced significantly more PKD-2::GFP tagged EVs than WT (A). (C) *mec-9(ok2853)* released significantly more PKD-2::GFP-tagged EVs. Significance determined by Mann-Whitney test; ***p<0.001 (D) Schematics of WT and *mec-9* mutant cephalic sensilla. In WT, EVs are released from cilium and are stored in lumen created by the sheath glial cell. Dashed lines (E-G) denote cross section level observed in images 4E, F, and G. *mec-9(ok2853)* mutants store and release excess EVs and contain dark and light matrix vesicles that contain EVs. (E-F) Cross section of the cephalic sensillum at the level of the transition zone in WT and *mec-9(ok2853)* males. CEM neurons shaded green, CEP neurons shaded blue. Scale bars 500 nm (E) Two EVs (arrows) observed in WT; one in the lumen and one in CEM cilium. (F) *mec-9(ok2853)* had a distended lumen with a significant increase of EVs and a lightly shaded matrix filled vesicle itself containing EVs. (G) Dark electron-dense matrix filled vesicles contained EVs. These vesicles were located at the level of the distal dendrite (See dotted line marked G in Figure 4D).

Next, we used transmission electron microscopy (TEM) to more closely examine the ultrastructure of the male cephalic organs containing EVs in ECM and lumenal spaces (Wang *et al*. 2014). In WT males, EVs are shed from the ciliary base of the CEM neuron and occupy a lumenal space formed by the glial support cells (Figure 4E). In *mec-9* mutant males, we observed three striking defects. First, in the *mec-9* cephalic lumen we observed a dramatic accumulation of EVs that ranged in diameter from 45nm to 226nm (Figure 4F). Second, we saw an increase in the volume of the glial lumen occupied by the EVs. The 2D area of the *mec-9* mutant lumen cross section was massively distended, perhaps due to EV hypersecretion and increased EV storage (Figure 4F). Eight WT and 12 *mec-9* cephalic sensilla were compared and revealed increased occurrences and larger diameters of lumenal spaces surrounding cilia and distal dendrites. The enlarged EV containing lumen is a phenotype observed in mutants that shed but do not environmentally release EV. Third, *mec-9* cephalic sensilla were filled with remarkable light and dark matrix filled vesicles, themselves also containing EVs. For example, the lightly shaded vesicle shown in Figure 4E was 506 nm in diameter and contained smaller vesicles ranging in diameter from 40-90 nm. Dark matrix vesicles found at the level of the distal CEM dendrite were 898 and 1056 nm in diameter and contained vesicles that ranged from 51-171 nm (Figure 4G). For both abnormal EV and matrix-filled vesicle phenotypes in *mec-9*, we observed larger complex vesicles in the glial cytoplasm as well as in the lumenal spaces surrounding the cilia, ciliary transition zone (TZ), and distal dendrite (data not shown). The presence of these larger vesicles with complex contents is a phenomenon not previously described in any of our other EV biogenesis mutants (Maguire *et al*. 2015; O’Hagan *et al*. 2017; Silva *et al*. 2017; Wang *et al*. 2014).. We propose that the *mec-9* and likely the *mec-5* extradendritic PKD-2::GFP ciliary localization (Cil) phenotype may correlate to extradendritic accumulation of the abnormal EVs in the ECM surrounding the CEM dendrite.

### *mec-9* and *pmk-1* act antagonistically in EV biogenesis and release

We wondered if the EV hypersecretion and excessive EV storage phenotypes seen in *mec-9* mutants were due to abnormal EV biogenesis or to defects in neuronal integrity. To test the former possibility, we examined genetic interactions with a positive regulator of EV shedding and release – the p38 MAPK *pmk-1*. We previously showed that *pmk-1* mutants do not accumulate EVs in the cephalic lumen and release fewer PKD-2::GFP labeled EV from ray neurons into the environment (Wang *et al*. 2015). In contrast, *mec-9* mutants exhibit the opposite phenotypes: hypersecretion and excessive release of cephalic EVs. WT and the *mec-9(ok2853); pmk-1(km25)* double mutant release comparable numbers of PKD-2::GFP labeled EVs (Figure 5A). In a *mec-9(ok2853); pmk-1(km25)* double mutant we see that EV release from ray neurons is restored to WT levels (Figure 5B), suggesting that *mec-9* and *pmk-1* may act antagonistically, with *mec-9* as a negative regulator, *pmk-1* as a positive regulator. Further, PKD-2::GFP fluorescence intensity (FI) and blind study of ciliary localization was analyzed and although the *mec-9* and *pmk-1* single mutants exhibit an increase of PKD-2::GFP at the ciliary base, the ciliary localization phenotype and increased ciliary FI is ameliorated in the double mutant (Figure 5C). We conclude that MEC-9 is a negative regulator of EV biogenesis, with the mutant shedding and releasing excessive amounts of EVs.

**Figure 5.**
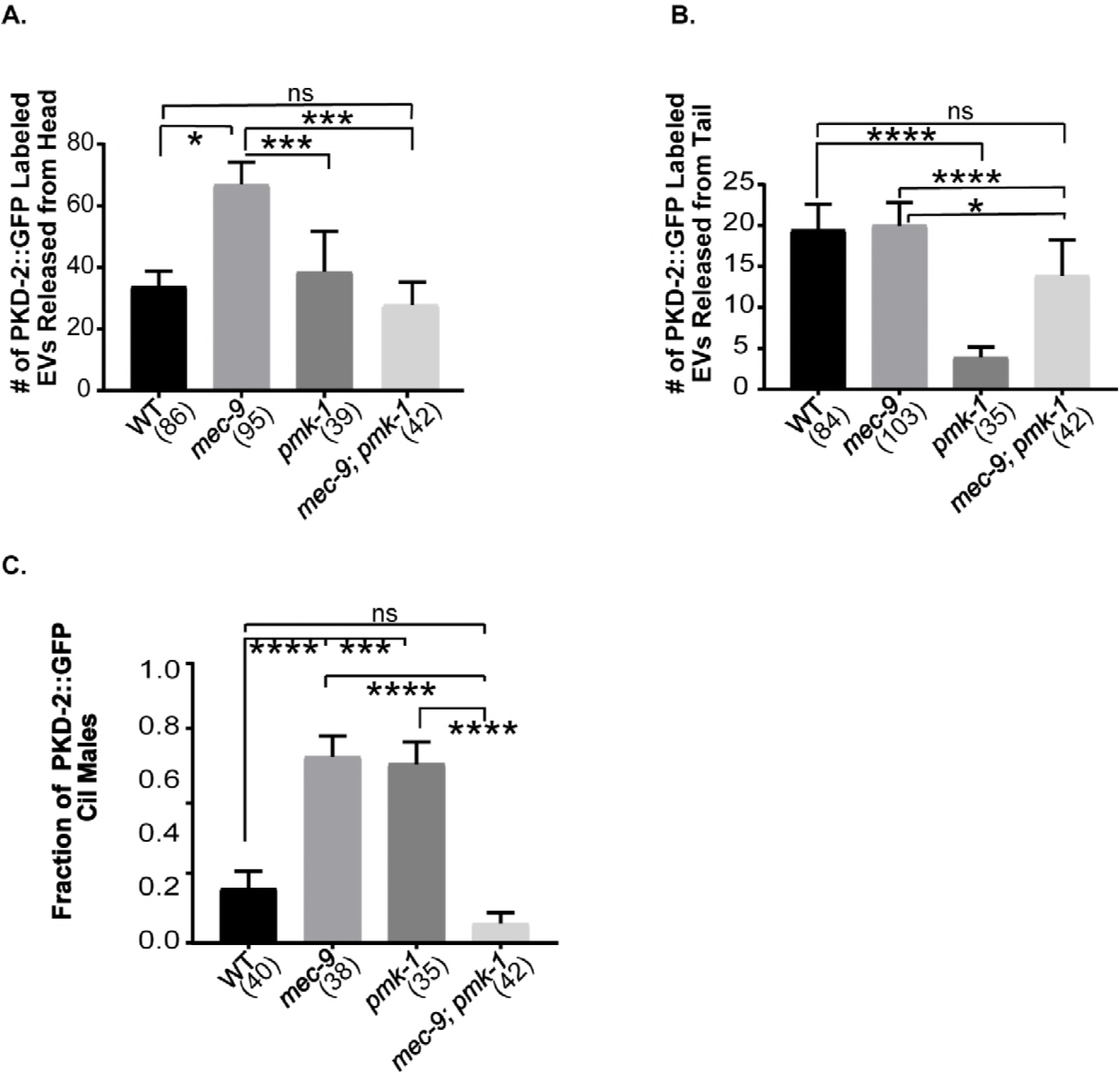
*mec-9* and *pmk-1* double mutant genetically suppressed single mutant EV biogenesis/release and PKD-2::GFP Cil phenotypes. (A) p38 MAPK *pmk-1(km25)* mutants are defective in RnB EV release (Wang et al., 2015). *pmk-1* mutation suppressed the *mec-9(ok2853)* abnormal EV hypersecretion in the head. (B) *mec-9* suppressed the *pmk-1* RnB EV release defect. (C) *pmk-1; mec-9* double mutant suppressed PKD-2::GFP Cil defect observed in both *mec-9* and *pmk-1* single mutant males. Significance was determined by Kruskal-Wallace test and Dunn’s multiple comparisons test. *p<0.05, **p<0.01, ***p<0.001, ****p<0.0001.

### *mec-9* maintains CEM and IL2 neuronal morphology

Given that *mec-9S* was expressed in many ciliated sensory neurons, we used transmission electron microscopy (TEM) to examine ultrastructure of sensory organs in the head of fixed age-matched young adult WT and *mec-9* males (Figure 6A). The head contains four cephalic sensilla, six inner labial sensilla, and two amphid sensilla, each of which performs distinct sensory functions in the worm (Inglis 2007).

**Figure 6.**
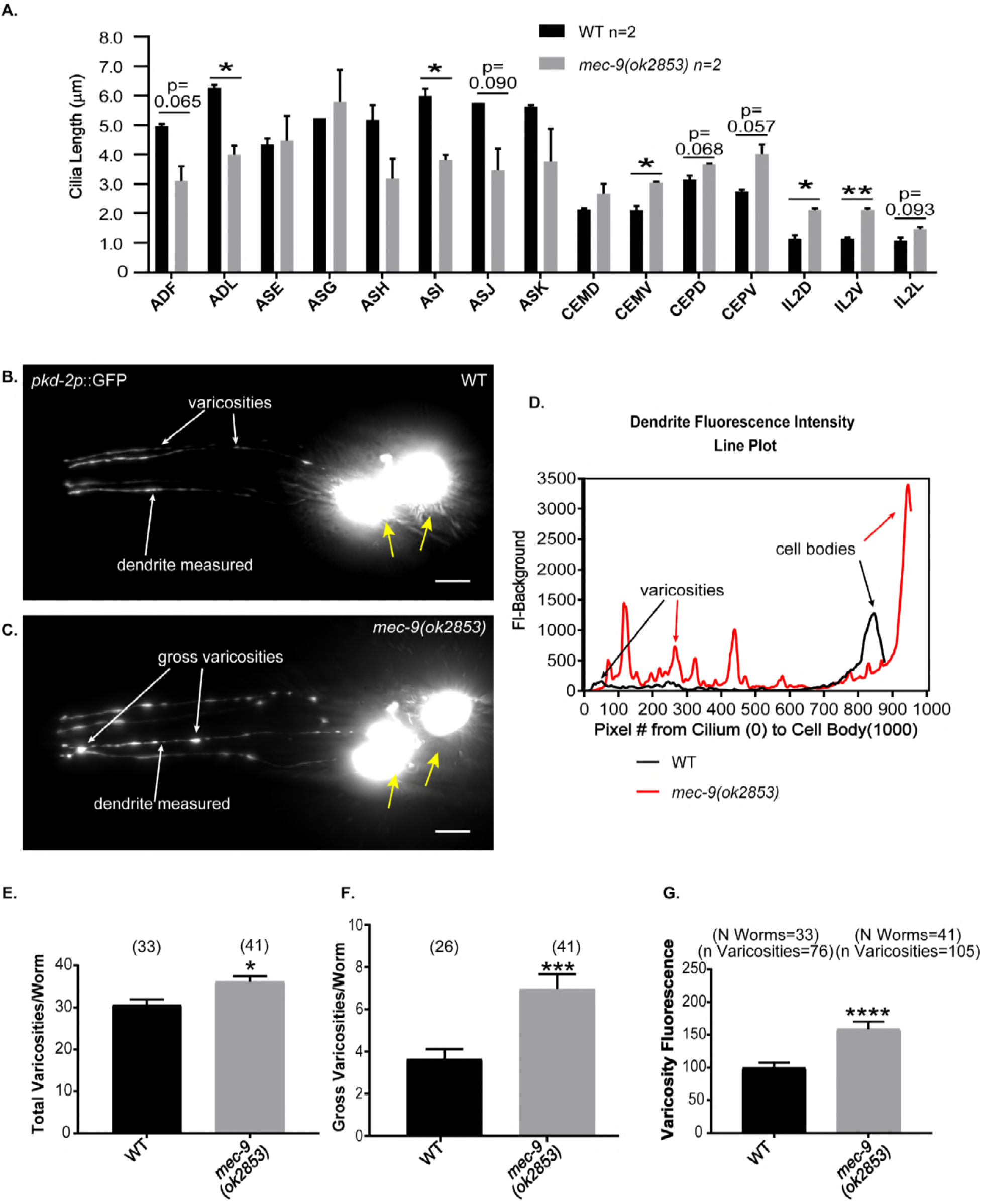
*mec-9* regulated neuron ciliary and dendritic anatomy. *mec-9* regulated CEM, IL2, and amphid channel ciliary lengths and maintained CEM neuronal integrity. (A) Amphid, double ciliated (ADL, ADF), amphid single ciliated (ASE, ASG, ASI, ASJ, ASK), cephalic (CEP) and male-specific CEM (ventral and dorsal), and inner labial (IL2) neurons were measured using serial sectioned electron micrographs (each layer~70 nm). Lengths were determined by counting number of 70 nm layers from transition zone to ciliary tip of each cilium. *mec-9(ok2853)* amphid cilia were shorter than WT and *mec-9(ok2853)* CEM, CEP and IL2 cilia were longer than WT cilia. Significance measured by Kruskal-Wallace test and Dunn comparisons test. *p<0.05, **p<0.01 (B-D.) *mec-9* maintained dendritic integrity. Images are maximum intensity Z-stacks of WT and *mec-9(ok2853)* males that expressed a *pkd-2p::GFP* transcriptional reporter. Images were brightened to observe dendritic morphology. Scale bars are 10 μm. Cell bodies denoted by yellow arrow. (B) Varicosities were observed in WT male CEMs, examples denoted by arrows. (C) *mec-9* mutant males have more varicosities. Large, “gross” varicosities denoted by arrows. (D) A line plot of WT (black) and *mec-9(ok2853)* (red) dendritic fluorescent intensity disclosed increased number of varicosities (E), increased gross varicosities (F), and that varicosities had a greater fluorescence. FI measured here using transcriptional reporter which allows for observation of neuron morphology only, not protein abundance measurements (G). Total varicosities and gross varicosities measured by blind study. Significance measured by Mann-Whitney test. Varicosity Fluorescence (G) normalized to 100. *p<0.05, ***p<0.001, ****p<0.0001.

In the male, the cephalic sensillum includes the cephalic male CEM neuron and the sex-shared CEP neuron. We viewed the CEM in cross section and measured from the ciliary transition zone to the ciliary tip and found WT CEM cilia to be ~2.1 μm in length (Figure 6A). In *mec-9* mutants, the CEM neurons were 1.2-1.5 times longer than WT. CEP cilia length was not significantly different between WT and *mec-9* mutant animals.

The inner labial sensilla house the IL1 and IL2 neurons. WT dorsal and ventral IL2 cilia averaged 1.155 μm and the lateral IL2s averaged 1.085 μm in length (Figure 6A). In the *mec-9(ok2853)* mutant, dorsal and ventral IL2 cilia were about twice as long as WT and the *mec-9(ok2853)* mutant lateral IL2s were ~1.3 times longer than WT (Figure 6A). IL1 ciliary lengths were similar in WT and *mec-9*, consistent with *mec-9S* expression in IL2 but not IL1 neurons.

The ECM encoding genes *mec-1, mec-5*, and *mec-9* regulate touch receptor neuron attachment and integrity (Pan *et al*. 2011). In *mec-5* and *mec-9* CEM neurons, PKD-2::GFP accumulated in extradendritic spaces (Figure 1 E,F, H). We therefore examined CEM neuronal morphology using a soluble GFP reporter. In WT and *mec-9* CEMs, *pkd-2p::GFP* was expressed and localized throughout the neuron in the cell body, dendrite, and cilium (Figure 6B). We did not observe extradendritic accumulation of soluble GFP in *mec-9* mutant, therefore the neurons are not fragile or leaky. However, *mec-9* mutant dendrites appeared brighter and more irregular than WT (Figure 6C).

To determine if there were differences in the dendritic morphology of WT and *mec-9* mutants, we compared FI peaks along the dendrite from the CEM cilium to cell body. These peaks represent dendritic varicosities and the example shown is indicative of the increased number and size of the varicosities in the *mec-9* mutant compared to WT (Figure 6D). WT animals averaged 30.6 varicosities per animal, whereas *mec-9* mutant animals averaged 36 varicosities (p= 0.0211) (Figure 6E).

We defined gross varicosities as those that are noticeably larger than normally observed and measure this phenomenon by blind investigation and FI. WT averaged 3.7 gross varicosities per animal and *mec-9* mutants averaged almost double (7 gross varicosities per animal) (p=0.0007; Figure 6F) The FI of the peaks/varicosities in WT were measured and normalized to 100. In comparison, *mec-9* mutant animals’ varicosities averaged 159% of WT intensity (p<0.0001) (Figure 6G). We conclude that the ECM gene *mec-9* was important for CEM dendritic morphology.

To determine if defects in dendritic morphology impacted protein transport in CEMs, we used time-lapse microscopy to measure the number and velocity of PKD::GFP particles moving along the dendrite. *mec-9(ok2853)* mutants had significantly fewer moving puncta (Table 1). In *mec-9(ok2853)*, PKD-2::GFP anterograde and retrograde velocities were significantly slower than WT (Table 1). Combined, these results indicate that the MEC-9 ECM component also regulated the CEM neuronal cytoskeleton and transport.

**Table 1.** *mec-9* regulated PKD-2::GFP particle abundance and velocity in CEM dendrites. The number and velocity of PKD::GFP particles was measured using time lapse fluorescent videos. TOTAL PARTICLES/μM: The average number of anterograde and retrograde PKD-2:GFP particles moving along the dendrite are reported here in particles/μm. *mec-9(ok2853)* mutants have significantly fewer particles than WT. ANTEROGRADE AND RETROGRADE VELOCITY: Anterograde and retrograde PKD-2::GFP particles velocities were measured in microns per second along the entire CEM dendrite. In *mec-9(ok2853)* males, PKD-2::GFP overall dendritic anterograde and retrograde particle velocity was slower than WT. Time lapse exposure per frame: 300 ms. Significance measured using Mann-Whitney test for particle number and Kruskal-Wallace test with Dunn’s comparison test for velocities.

| | MEAN ± SEM | n | N | P VALUE | DIFF. FROM WT |
| --- | --- | --- | --- | --- | --- |
| **TOTAL PARTICLES/μM** |  |  |  |  |  |
| WT | 82.66 ± 5.199 | 48 | 18 | | |
| *mec-9(ok2853)* | 63.97 ± 2.976 | 83 | 23 | 0.0241 | -18.68 ± 5.559 |
| **ANTEROGRADE VELOCITY** |  |  |  |  |  |
| WT | 0.8215 ± 0.007 | 4654 | 18 | | |
| *mec-9(ok2853)* | 0.7496 ± 0.005 | 4765 | 23 | <0.0001 | -0.07186 ± 0.009 |
| **RETROGRADE VELOCITY** |  |  |  |  |  |
| WT | -0.7607 ± 0.006 | 4971 | 18 | | |
| *mec-9(ok2853)* | -0.6551 ± 0.004 | 5019 | 23 | <0.0001 | -0.1056 ± 0.008 |

### *mec-9* regulates amphid ciliary length, positioning, and fasciculation

*mec-9S* was also expressed in the dye-filling amphid sensory neurons that are housed in bilateral amphid sensilla (Figure 2H-I). Dye filling assays assess ciliary integrity: WT animals dye fill whereas ciliary mutants are dye filling defective (Dyf). *mec-1(e1066)* animals have a slight Dyf phenotype (Perkins *et al*. 1986). We performed dye filling assays to determine whether *mec-5* and *mec-9* have similar ciliary defects as seen in *mec-1. mec-1(e1066), mec-5(e1503)*, and *mec-9(ok2853)* had subtle amphid Dyf defects with 1-4 cells of 12 not filling (Figure 7A). *mec-1(e1066)* and *mec-5(e1503)* phasmids are also Dyf, with 1-2 out of four cells not filling (Supplemental Figure 5). Therefore, the amphid and phasmid ciliated sensory neurons also require proteins encoded by *mec-1, mec-5*, and *mec-9* ECM genes.

**Figure 7.**
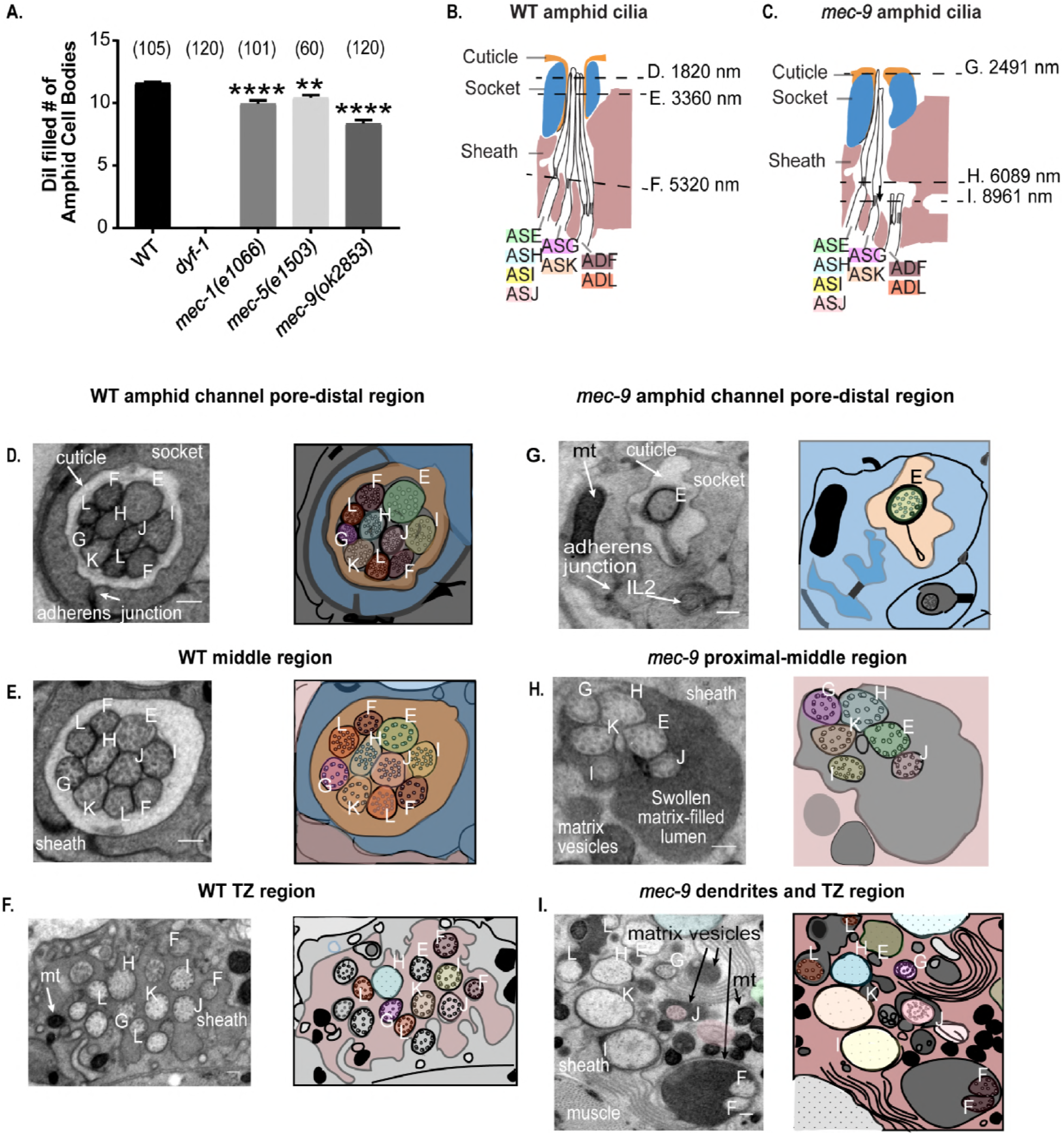
*mec-1, mec-5*, and *mec-9* mutants are dye filling defective and *mec-9* is necessary for amphid cilia organization and ECM deposition. (A) Number of amphid neurons out of 12 that filled with dye. (B-C) Schematics of WT and *mec-9(ok853)* amphid sensory organ (not all neurons shown). Proximal and medial sections of WT amphid cilia were in the sheath. Distal cilia were surrounded by socket and inverted cuticle with cilia tips exposed to the environment. Dashed lines denote levels of cross sections shown in (D-I). (C) *mec-9(ok2853)* amphid cilia were short and displaced with few ciliary tips exposed to the environment. (D) Left amphid channel pore and its schematic showing ten cilia at level of singlet microtubules. WT double ciliated amphids were found dorsally and ventrally in the pore with the ADF (medial) and ADL (lateral). The WT single ciliated amphids are found in the center two rows of amphids. The top row of single ciliated amphid channel neurons from medial to lateral are ASE, ASH, and ASG. The bottom row: ASI, ASJ, and ASK. WT left and right (not shown) amphids are mirror images of each other. Neurons are labeled by final amphid letter. (E) WT middle segment contained 10 cilia with microtubules arranged concentrically. (F) We observed the transition zones of most WT amphid cilia at this level and they were embedded within the amphid sheath. (G) Only one cilium out of ten was visible in *mec-9* mutant channel pore. Socket and sheath had abnormal morphology as compared to WT. (H) *mec-9* mutant middle segment: six out of ten cilia were visible in the lumen and there was increased matrix. *mec-9* mutant amphids were abnormally arranged in three rows of two cilia. The top row: ASG and ASH, the second row: ASK and ASE, and the bottom row: ASI and ASJ, all of which are single ciliated amphids. Increased numbers of matrix filled vesicles were observed. (I) The double ciliated amphids (ADFs and ADLs) were in adjacent, aberrant electron dense spaces (increased matrix). Scale bar 200 nm.

We proceeded to examine amphid sensilla ultrastructure using TEM. In WT, 10 amphid cilia are enclosed in a pore of the bilateral papilla at the worm nose and surrounded by the cuticle (Figures 7B and 7D). *mec-9* worms exhibited a misshapen cuticular pore encapsulating only the ASE cilium (Figure 7C). In the WT amphid channel, ten cilia are present in an invariant and precise arrangement (Figures 7D and 7G). In *mec-9* mutants, we observed only cilia from the ASE, ASG, ASH, ASI, ASJ, and ASK (the single ciliated amphids) in the more proximal amphid channel; ADF and ADL (the double ciliated neurons) were displaced in an adjacent, aberrant electron dense space (Figure 7I). Consistent with a function in these neurons, *mec-9shp::GFP* was expressed in ADL and ADF. We conclude that *mec-9* was necessary for cilia alignment and placement in amphid sensilla.

We measured ciliary lengths in these double ciliated amphid neurons from the bifurcation at the transition zone to ciliary tip. In contrast to the longer CEM and IL2, EV releasing cilia in *mec-9* mutants, the cilia of the ADL and ADF amphid neurons were significantly shorter than WT by ~1.5x (Figure 6A). We also measured single ciliated amphid axonemes and found the ASI cilia significantly shorter than WT (Figure 6A). These amphid cilia length defects may contribute to dye filling defects observed in *mec-9* mutants (Figure 7A). Not only were *mec-9* amphid cilia shorter and misplaced, *mec-9* mutant amphid transition zones were disorganized and abnormally dispersed along the anterior-posterior axis (compare WT in Figure 7F to *mec-9* in Figure 7I), suggesting that the *mec-9* ECM component is necessary for regulating dendritic fasciculation and/or transition zone placement.

Perkins et al. describe the presence of dark matrix vesicles in the space between the amphid cilia and sheath in WT animals and highlighted several mutants (*osm-1* and *osm-3*) that abnormally accumulate large matrix vesicles (Perkins *et al*. 1986). In *mec-9(ok2853)* mutants, we observed a substantial increase in electron dense extracellular spaces that we identified as large matrix filled vesicles (Figure 7H). *mec-9* mutants also had excessive matrix filled spaces in the amphid neuron sheath surrounding the amphid neurons that decouples cilia from glia (Figures 7H). Because of these spaces, there is a dysmorphic amphid socket and sheath in *mec-9* mutant animals, which may contribute to the disorganization of the amphid channel cilia (Figures 7G-I). We conclude that *mec-9* regulates ECM deposition in amphid and cephalic sensilla (Figure 4H-I).

## DISCUSSION

ECM is important for neuronal anatomy and organization of the brain and nervous system (Gardiner 2011). For example, the ECM glycoprotein Reelin is necessary for migration of neocortical radial cells in the mammalian brain (Franco *et al*. 2011). In *C. elegans*, ECM components *dex-1* and *dyf-7* regulate amphid dendritic extension, which then affects cilia placement (Heiman and Shaham 2009). Mutations in the RNA splicing gene *mec-8* or ECM component *mec-1* mutants cause ciliary fasciculation defects (Perkins *et al*. 1986). The ECM gene *mec-9* is expressed in the *C. elegans* nervous system where it provides mechanical support to multiple cell types including non-ciliated touch receptor neurons (Du *et al*. 1996) and, as shown here, ciliated sensory neurons.

*mec-9* encodes two isoforms (Du *et al*. 1996). The *mec-9* long isoform is expressed in touch receptor neurons (Du *et al*. 1996). Here, we show that the *mec-9* short isoform is expressed in the shared and male-specific ciliated nervous system. In amphid and phasmid sensory organs, *mec-9* mutants are dye filling defective, which reflects abnormalities in stereotypical ciliary positioning and fasciculation. In cephalic sensory organs of males, *mec-9* mutants display distended extracellular spaces that contain excessive amounts of ciliary EVs and longer cilia. In CEM neurons, *mec-9* is also required for dendritic integrity, with *mec-9* mutant dendrites showing varicosities and waves as opposed to more linear trajectories. This abnormal dendritic morphology is typically observed in the nervous system of aged animals, and these age-dependent defects are accelerated by mutations that disrupt neuronal excitability or mechanosensation, including the *mec-1, mec-5*, and *mec-9* ECM genes (Pan *et al*. 2011). Further studies are needed to ascertain if MEC-9 protein physically restrains the dendrite or attaches to the ciliary or dendrite membrane to properly position the neuron. However, it is clear that MEC-9 and the ECM support the development and function of the *C. elegans* sensory nervous system (Figure 8).

**Figure 8.**
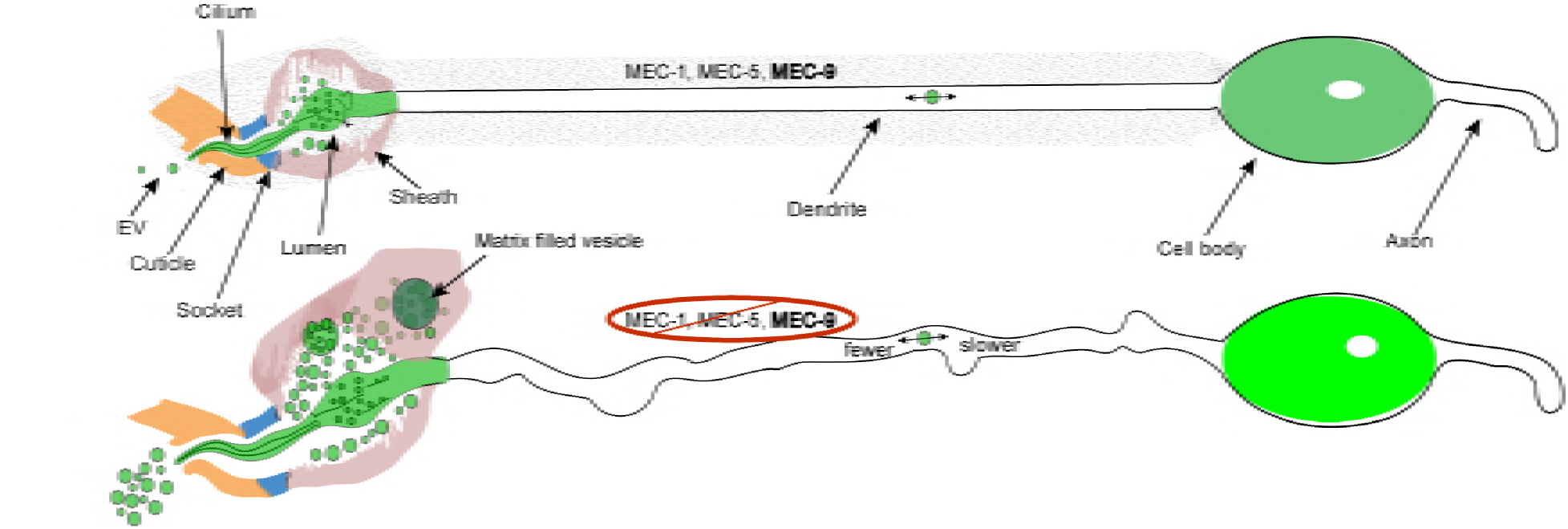
MEC-1, MEC-5, and MEC-9 ECM influence ciliated neuronal structure, neuronal transport, and neuronal function. The ECM genes *mec-1, mec-5*, and *mec-9* regulate PKD-2::GFP localization and male mating behaviors (Figure 1 and 2). *mec-9* also is necessary for PKD-2::GFP dendritic transport (Table 1), negative regulation of EV biogenesis, storage, and release (Figure 4), and neuron anatomy, such as dendritic integrity (Figure 6), cilia length (Figure 6) and organization (Figure 7), and matrix deposition (Figures 4G and 7H-I).

ECM and cilia share an intimate association. In aortic valves, primary cilia are restricted to ECM zones (Toomer *et al*. 2017). Chondrocytes have cilia embedded in ECM (Ruhlen and Marberry 2014). In umbilical cord mesenchyme, ECM regulates ciliary orientation (Nandadasa *et al*. 2015). ECM regulates ciliogenesis and organogenesis of Kupffer’s vesicle, the zebrafish equivalent of the mammalian embryonic node (Compagnon *et al*. 2014; Hochgreb-Hagele *et al*. 2013). In Drosophila, the ECM protein artichoke is required for morphogenesis of ciliated organs (Andres *et al*. 2014). The ECM gene spacemaker/Eyes shut/RP25 is necessary for ciliary pocket morphology and photoreceptor survival (Yu *et al*. 2016), with mutation causing photoreceptor degeneration in retinitis pigmentosa patients (Abd El-Aziz *et al*. 2008).

ECM influences ciliary length in multicellular animals. Here we show that ECM regulates ciliary length in a cell-type specific manner in *C. elegans*. In amphid channel neurons, *mec-9* is a positive regulator, with *mec-9* mutants having shorter cilia. Conversely, in EV-releasing IL2 and CEM neurons, *mec-9* is a negative regulator of ciliary length, with *mec-9* mutants having significantly longer cilia. In mammalian skin, ECM component laminin-511 and its receptor integrin-b1 are required for primary cilia formation (Gao *et al*. 2008). In mouse embryonic fibroblast 3T3-L1 cells, type 1 collagen promotes primary ciliary growth by repressing the HDAC6-autophagy pathway (Xu *et al*. 2018). In zebrafish Kupffers Vesicle, laminin-1 is a positive regulator of ciliary length (Compagnon *et al*. 2014; Hochgreb-Hagele *et al*. 2013). The mechanisms by which ECM controls ciliary length are largely unknown. A positive or negative feedback loop may act cell autonomously (between the ciliated cell and ECM secreted by the cell itself) or non-autonomously (between the ciliated cell and ECM secreted by neighboring cells). Our data are consistent with both possibilities. Rescue experiments indicate that *mec-5* acts non-autonomously while *mec-9S* expression in ciliated neurons suggest cell autonomous function.

EVs are components of the ECM and EVs themselves may carry ECM proteins as cargo (Rackov *et al*. 2018; Rilla *et al*. 2017). Chlamydomonas ciliary EVs carry ECM proteins and ECM-degrading proteases, including a proteolytic enzyme that degrades ECM necessary for hatching (Long *et al*. 2016; Wood *et al*. 2013). Here we show that *mec-9* mutants display dramatic increases in EV shedding and release (Figure 4C) and abnormal dark/light matrix vesicles containing EVs (Figure 4G), suggesting that the MEC-9 ECM component negatively regulates both EV shedding and release. We do not understand how ECM regulates EV biogenesis. However, genetic analysis revealed that *mec-9* and the p38 MAPK *pmk-1* acted antagonistically in EV biogenesis and release. *pmk-1* mutants were defective in EV shedding and EV release (Wang *et al*. 2015). Interestingly, *pmk-1* suppressed the *mec-9* EV hypersecretion phenotype and *mec-9* suppressed *pmk-1* EV hyposecretion (Figure 5), suggesting that these genes act in opposing pathways that control EV biogenesis. An intriguing possibility is that MEC-9/ECM and *pmk-1* kinase regulate the same target(s) such as a cell surface ECM receptor. In mice, CELSR3 (cadherin EGF LAG seven-pass G-type receptor 3) has MEC-9-like EGF domains in its N-terminal ectodomain and CELSR3 interacts with a kinase that regulates extension and guidance of sensory neurons (Goffinet and Tissir 2017). Mice deficient in CELSR2 and CELSR3 are defective in ependymal cilia development and develop hydrocephalus, a ciliopathy phenotype (Goffinet and Tissir 2017). The ligand(s) that activates CELSR2 and CELSR3 are not known.

The CELSR family is categorized as adhesion GPCRs (Liebscher and Schoneberg 2016). Adhesion GPCRs (aGPCRs) contain a large N-terminal ectodomain that contains a tetherized agonist *Stachel* sequence (Liebscher and Schoneberg 2016). Removal or structural changes to the N-terminal ectodomain exposes the *Stachel* sequence, which in turn activates the GPCR. Proposed mechanisms of Stachel release and activation of aGPCR signaling include mechanical stress and binding of ECM proteins to the N-terminus (Liebscher *et al*. 2014; Luo *et al*. 2014; Scholz *et al*. 2016).

The polycystin-1 family, while an 11-transmembrane spanning receptor class, has some features similar to an aGPCR (Cazorla-Vazquez and Engel 2018; Langenhan *et al*. 2015; Trudel *et al*. 2016). The function of the polycystins remains an enigma, even thirty years after the cloning of PKD1 and PKD2 (Ma *et al*. 2017). Based on their ciliary localization, the polycystins were thought to be ciliary mechanosensors, but this model was disproven by the Clapham lab {Delling, 2013 *#*14868}(DeCaen *et al*. 2013). In mice, an ECM receptor integrin signaling pathway is essential for the development of ADPKD (Lee *et al*. 2015). An intriguing possibility is that the ECM itself acts in permissive fashion to allow cell-type specific polycystin activation and signaling, a mechanism used by adhesion GPCRs. Several lines of evidence support this idea. - Polycystin-1 binds to many ECM proteins including collagen I, II, and IV, fibronectin, and laminin (Malhas *et al*. 2002). Inactivation of integrin-b1 or integrin-linked kinase inhibits cystogenesis in *Pkd1* mutant mice (Lee *et al*. 2015; Raman *et al*. 2017). In zebrafish, pkd2 deficiency causes increased collagen synthesis via upregulated protein secretion and downregulation of this secretory pathway rescues cystogenesis (Le Corre *et al*. 2014; Mangos *et al*. 2010). Combined, these studies reflect the close but poorly understood association between ECM, cilia, and the polycystins.

Our data reveal the profound importance of ECM components in nervous system of the worm. Model Figure 8 depicts the activity of ECM genes on ciliated sensory neurons. *mec-1, mec-5*, and *mec-9* regulated PKD-2::GFP localization and male mating behaviors. *mec-9* also influenced PKD-2::GFP dendritic transport and negatively regulated EV biogenesis, storage, and release. *mec-9* and ECM was also important for neuronal anatomy, dendritic integrity, ciliary length and organization, and matrix deposition. Notably, abnormal ECM is implicated in the pathogenesis of ADPKD (Calvet 1993; Song *et al*. 2017) and the polycystins interact with ECM and focal adhesion proteins (Drummond 2011; Retailleau and Duprat 2014). Renal fibrosis observed in PKD is characterized by excessive deposition of ECM proteins (Drummond 2011; Song *et al*. 2017). Here we show that ECM is necessary for the health and well-being of ciliated neurons and neural organs in the nematode. Our findings highlights the promiscuity of ECM components, reveal ECM activity in ciliated neurons of the worm, and broadens the scope of activity of the ECM proteins originally named for their roles in mechanosensory touch receptor neurons. That ECM proteins contribute to ciliary localization and function of the polycystins in *C. elegans* advances the understanding of ciliopathies like ADPKD.

## Acknowledgements

We thank Brian Coblitz and Martin Chalfie (Columbia University) for *mec-5(+)* rescue constructs; Richard Poole (Hobert lab, Columbia University) for assistance with whole genome sequencing of *cil-7/mec-5(my2);* Aranzta Barrios for advice on leaving assays and statistical analysis; Geoff Perumal and Leslie Gunther-Cummins for their help with HPF/FS processing; Monica Driscoll, Marion Gordon, Sunita Kramer, Barth Grant, Martha Soto, Christopher Rongo, and the Rutgers Super Worm Group for essential feedback during D.D.’s graduate career; the Barr lab for ongoing discussion and constructive criticisms, and especially Robert O’Hagan for his expertise in mechanotransduction; Gloria Androwski for outstanding laboratory support; WormBase and WormAtlas for online resources. This work was funded by National Institutes of Health grants DK059418 and DK111214 (to M. M. B.), OD 010943 (to D. H. H), and F31DK103550 (to D.D.). Some strains were provided by the National BioResource Project and the *Caenorhabditis* Genetics Center (CGC), which is funded by NIH Office of Research Infrastructure Programs [P40 OD010440]. Authors declare no competing financial interests or any funding that can compromise the integrity of this work.

## Supplemental Figures

**Table S1.** *mec-1, mec-5* and *mec-9* mutants regulate PKD-2::GFP abundance in male specific neurons of the head (CEM) and tail (RnB). CEM and RnB ratio (cilium/cell body or part of neuron specified) of maximum or mean fluorescence intensity (as denoted in table) showed that PKD-2::GFP abundance in *mec-1, mec-5* and *mec-9* mutants was variable. For example (cilium/cell body) of maximum intensity showed that PKD-2::GFP abundance in CEM cilia is increased in comparison with the cell bodies only in *mec-9* ECM gene mutants (Figure 1J); however, mec-1 and *mec-5* alleles also affected PKD-2 abundance. Background measurements were subtracted from cilium and cell body values for standardization of images and we expressed the measurements in ratio of cilia to cell body FI. Significance was measured by Kruskal-Wallace test, comparisons made using Dunn’s multiple comparisons. Wild type animal values were normalized to 100. The *mec-9* mutants had the brightest maximum FI when compared than WT *p<0.05, **p<0.01, ***p<0.001, ****p<0.0001.

|  | **WT n=54** | **Cil=cilium**<br>**CB=cell body**<br>**Den=dendrite**<br>**BG=background** | ***mec-1***<br>**(*e1066*)**<br>**n=72** | ***mec-5***<br>**(*e1503*)**<br>**n=31** | ***mec-9***<br>**(*ok2853*)**<br>**n=59** |
| --- | --- | --- | --- | --- | --- |
| **CEM**<br>**(head)** | **cilium/cell body** | **Cil/CB mean** | ↓ * | ns | ↑ **** |
|  |  | **Cil/CB max** | ↓ * | ns | ↑ ** |
|  |  | **Den/CB mean** | ↓ * | ↑ * | ↓ **** |
|  |  | **Den/CB max** | ↑ * | ↑ ** | ns |
|  | **FI-background** | **Cil-BG mean** | ns | ns | ↑ **** |
|  |  | **Cil-BG max** | ns | ns | ↑ **** |
|  |  | **Den-BG mean** | ns | ns | ↑ **** |
|  |  | **Den-BG max** | ↑ *** | ns | ↓ **** |
|  |  | **CB-BG mean** | ↑ * | ↓ * | ↑ **** |
|  |  | **CB-BG max** | ns | ↓ **** | ↑ **** |
| **RnB**<br>**(tail)** | **cilium/cell body** | **Cil/CB mean** | ns | ns | ns |
|  |  | **Cil/CB max** | ↓ ** | ns | ↑ * |
|  |  | **Den/CB mean** | ↓ * | ↑ * | ↓ * |
|  |  | **Den/CB max** | ns | ns | ns |
|  | **FI-background** | **Cil-BG mean** | ns | ns | ↑ * |
|  |  | **Cil-BG max** | ↓ * | ↓ **** | ↑ *** |
|  |  | **Den-BG mean** | ns | ns | ns |
|  |  | **Den-BG max** | ns | ns | ↑ ** |
|  |  | **CB-BG mean** | ns | ns | ↑ * |
|  |  | **CB-BG max** | ns | ns | ↑ ** |

**Figure S1.**
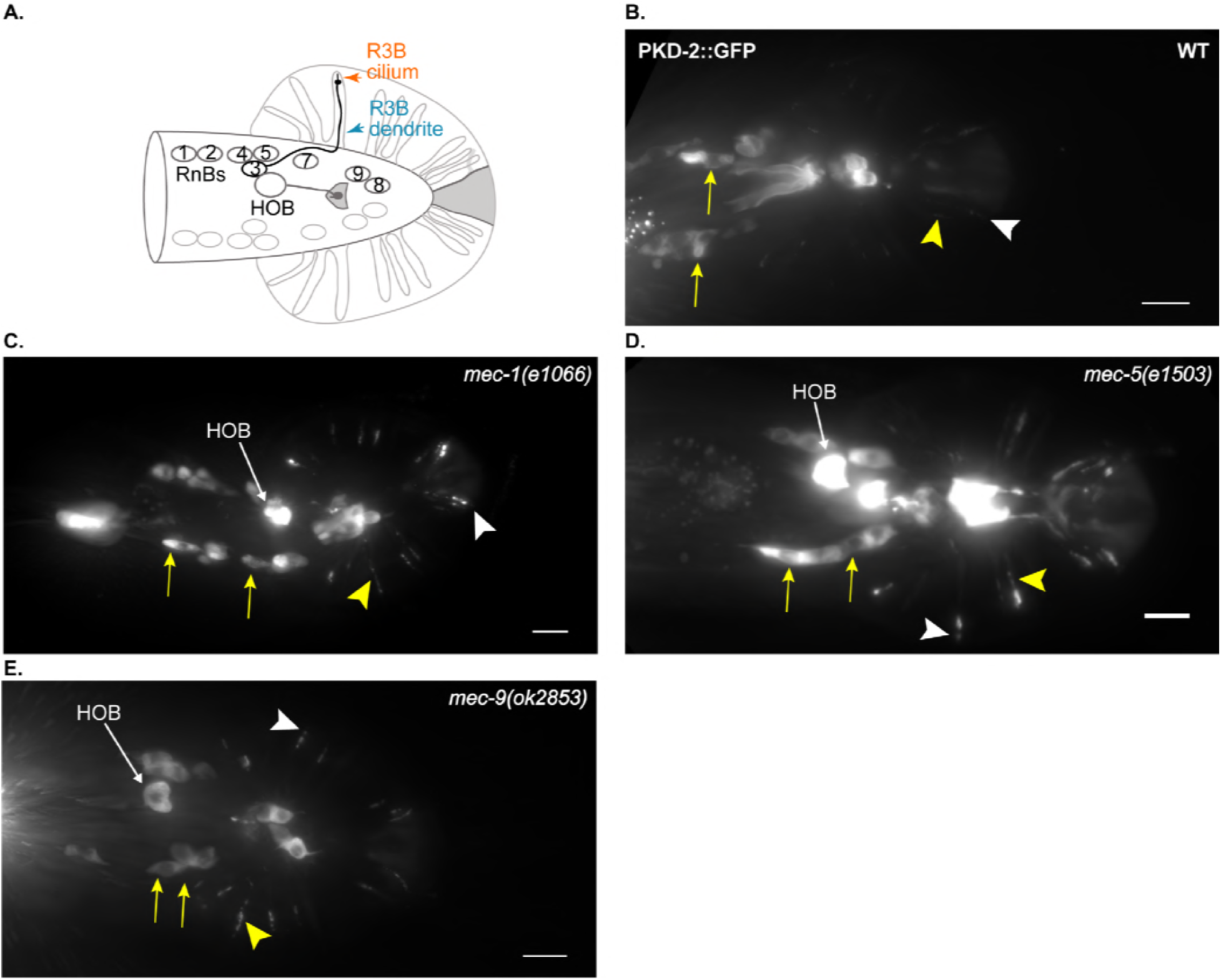
Alleles of *mec-1, mec-5* and *mec-9* regulate PKD-2::GFP localization and abundance in RnBs. A. Schematic of WT RnBs and HoB male neurons in the tail of a male (green). (B-E) Images were compiled from a 630x maximum intensity Z-series obtained by fluorescent microscope. Scale bar is 10 μm. White arrow head, cilium; cell body, yellow arrow; dendrite, yellow arrowhead. (B) A ventral view of WT PKD-2::GFP (translational reporter). GFP localized to RnB and HOB cell bodies and cilia. (C) Dorsal view of *mec-1(e1066)* here showed increased PKD-2::GFP at RnB cilia and ciliary base but statistically there was decreased FI (Supplemental Table 1). (D) *mec-5(e1503)* PKD-2::GFP ciliary localization showed increased PKD-2::GFP at RnB cilia and ciliary base but overall there was no statistical difference from WT (Supplemental Table 1). (E) We observed increased PKD-2::GFP at *mec-9(ok2853)* at RnB cilia and ciliary base and a significant increase in FI (Supplemental Table 1).

**Figure S2.**
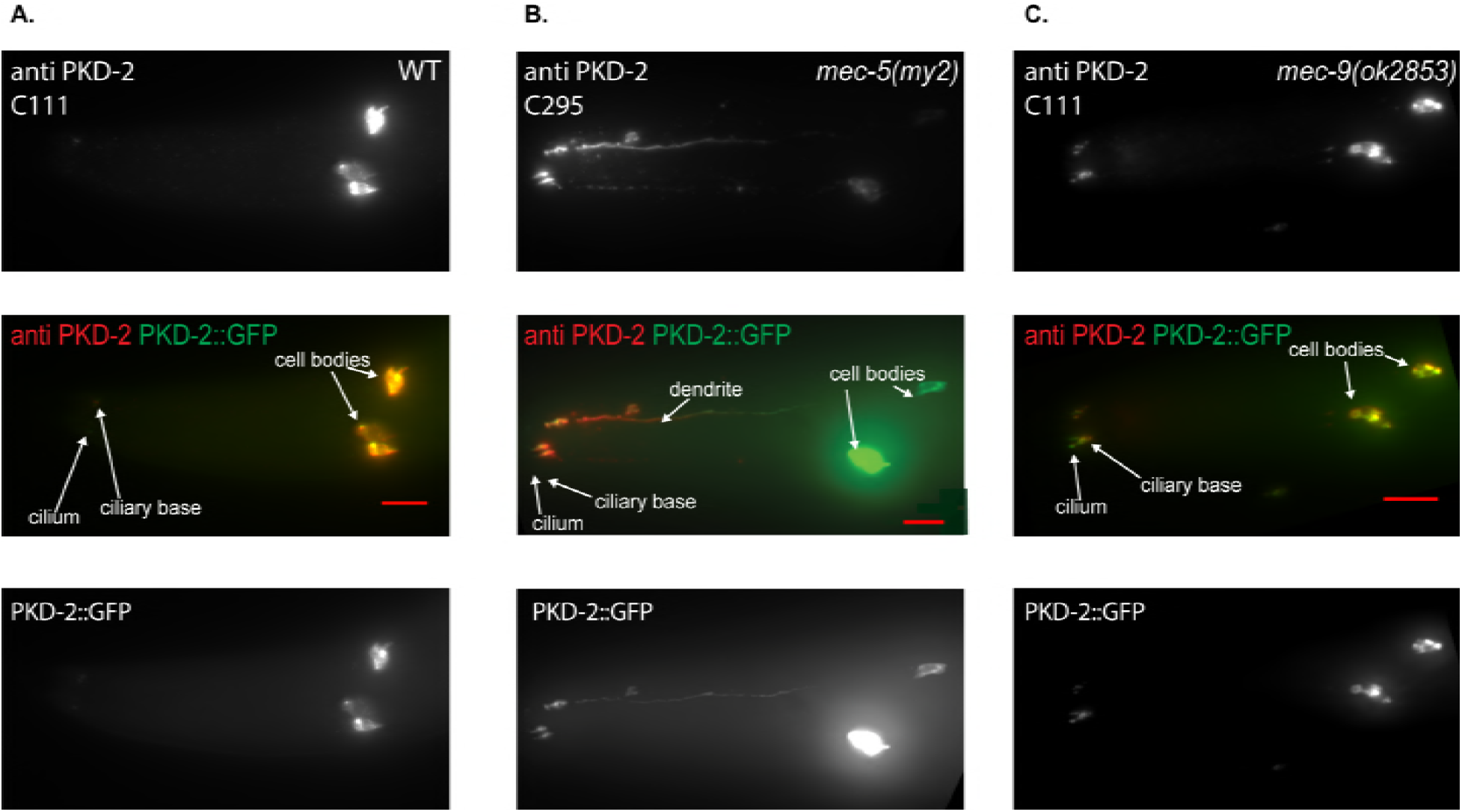
Anti-PKD-2 antibody stained CEM neurons. Antibody fluoresces with PKD-2::GFP in CEM neuron. Anti-PKD-2 in *mec-5(my2)* males have characteristic extradendritic accumulation seen in both antibody and PKD-2::GFP. Shown here are two different antibodies. Clone C111 is generated to an epitope at the N-terminus of PKD-2. This PKD-2 antibody aggregated in the cell body of WT (A) and *mec-9(ok2853)* (B). Clone C295, was also generated to an epitope at the N-terminus. C295 is a non-aggregating PKD-2 antibody that was used in *mec-5(my2)* (B). We observed no overt differences in PKD expression or intensity in *mec-9(ok2853)* when anti-PKD-2 and PKD-2::GFP are compared. Scale bar is 10 μm.

**Figure S3.**
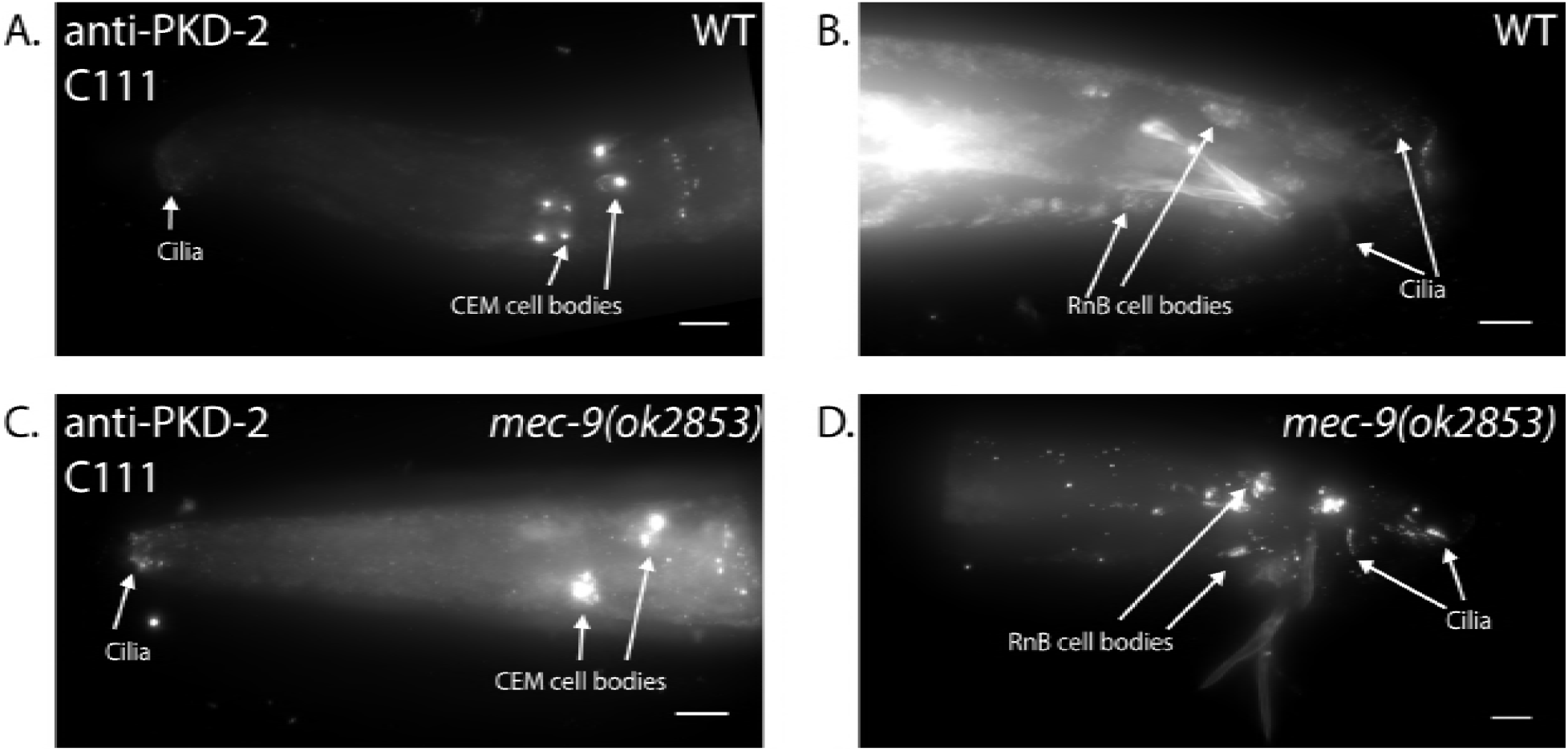
N-terminal anti-PKD-2 (C111) antibody showed increased, endogenous localization at male specific CEM and RnB cilia. In WT, endogenous PKD-2 was limited to the cell bodies and cilia of CEM head neurons (A) and ray RnB and hook HOB tail neurons (B). *mec-9(ok2853)* males are Cil (C) and endogenous PKD-2 mislocalized to dendrites and (D). Scale bar 10 μm

**Figure S4.**
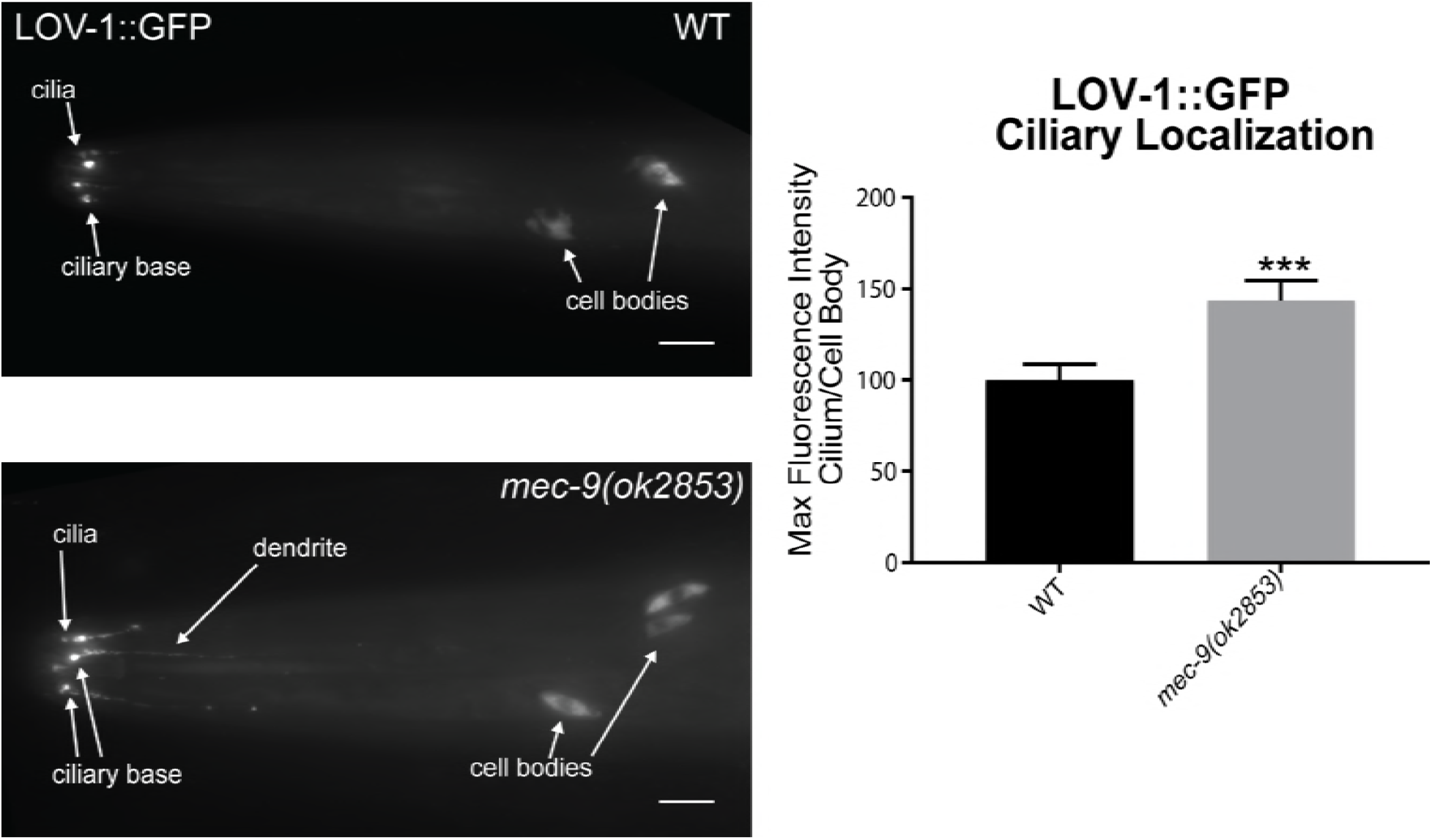
MEC-9 regulates LOV-1::GFP ciliary localization. LOV-1::GFP mislocalized to CEM distal dendrites in mutant animals. Scale bar is 10 μm. Quantitative measurement of fluorescence intensity in *mec-9* mutants revealed an abundance of LOV-1::GFP at the cilia and cilium proper of male specific CEM neurons when in ratio to CEM cell body. Significance measured by Mann-Whitney test. **p<0.01

**Figure S5.**
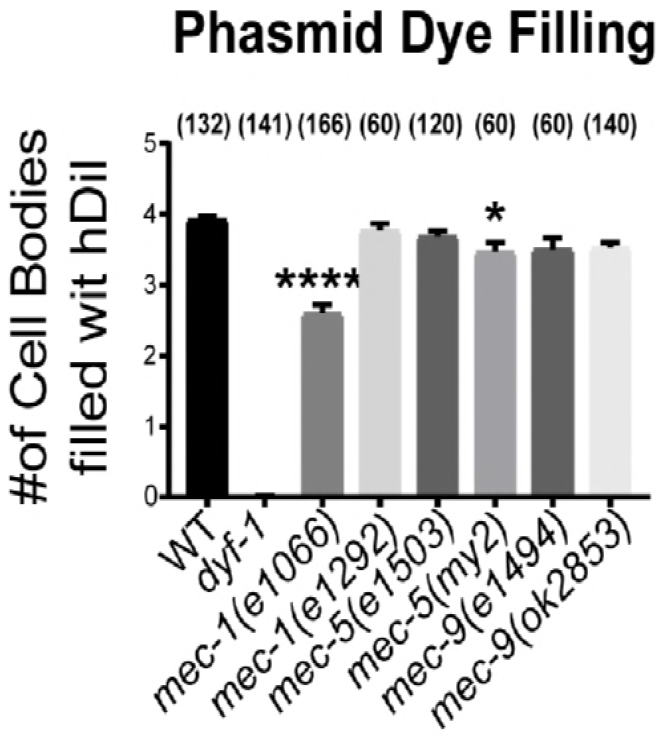
*mec-1(e1066)* and *mec-5(e1503)* phasmids are Dyf. In *mec-1(e1066)* and *mec-5(e1503)*, one-two out of four phasmid cells did not fill. Significance measured by Kruskal-Wallace test with Dunn comparisons. ****=<0.0001, *p<0.05.

